# Cortical ER-PM contact sites promote nuclear ER membrane integrity and mitotic nuclear segregation

**DOI:** 10.64898/2026.11.09.765494

**Authors:** Aleksa Nenadić, Jason R.B. Gurung, Matthew W. Volpiana, Valentin Jaumouillé, Christopher T. Beh

## Abstract

The endoplasmic reticulum (ER) extends from the nuclear envelope through the cytoplasm to the cell cortex, where it contacts the plasma membrane (PM) at ER-PM membrane contact sites (MCSs). We hypothesized that as contiguous extensions of nuclear ER these cortical ER (cER) attachments to the PM might dampen shearing stresses on the nuclear envelope and help orchestrate nuclear ER dynamics. Although individual cER-PM tether proteins promote cER inheritance during mitosis, our findings reveal that removing all cER-PM tethers collectively disrupts nuclear envelope membrane integrity. In yeast Δ-super-tether (Δ-s-tether) cells lacking seven cER-PM tethers, cER detachment causes membrane “blisters” and nuclear envelope ruptures; however, individual tether deletions or ER stress do not disrupt nuclear containment. Despite the extensive nuclear envelope lesions in Δ-s-tether cells, ESCRT-III (Endosomal Sorting Complex Required for Transport-III)- mediated membrane repair sustains cell viability. The loss of nuclear integrity, however, appears to disrupt genomic repair and promotes spontaneous mutation; separately, nuclear inheritance is also disrupted. Without cER-PM MCSs, cortical actin depolarization and defects in nucleopodia impair mitotic nuclear dynamics and segregation, which triggers a Spindle Position Checkpoint (SPOC) cell-cycle delay. That cER-PM MCSs serve as dual-function hubs for the preservation of nuclear envelope integrity is indicated by two independent mechanisms capable of rescuing the observed nuclear membrane defects: the restoration of phospholipid metabolism or the re-establishment of nonspecific cER-PM contact.

## Introduction

Dysregulation of endoplasmic reticulum (ER) membrane structure, in its various forms, can lead to cancer, metabolic disorders, and neurodegenerative diseases [1–7]. In its entirety, the ER is organized as a contiguous network of sheets and tubules extending from the nuclear envelope, through the cytoplasm, to the cell cortex, where it associates directly with the plasma membrane (PM) [8–10]. In yeast, almost half of the inner surface area of the PM is coated with cortical ER (cER) attached at specific sites where the two membranes in direct contact. cER-PM membrane contact sites (MCSs) are established by protein tethers that maintain the cER in close proximity (∼10-50 nm) to the inner PM surface, allowing rapid exchange of metabolites between the two membranes [11–13]. In this regard, cER-PM MCSs represent an important nexus for coordinating regulatory interactions between these two membranes. Because all ER membranes are contiguous, including that surrounding the nucleus, we hypothesize that cER-PM MCSs affect the ER at the cell cortex, throughout the cytoplasm, as well as the nuclear ER.

Under standard growth conditions, contact between the cER and PM is mediated by seven primary ER tethering factors, the elimination of all abolishes directed cER attachment to the PM [14]. In yeast Δ-super- tether (Δ-s-tether) cells, which lack seven cER-PM tethering proteins, cortical ER is almost entirely absent, and normal phospholipid regulation and lipid metabolism is disrupted [14]. As revealed through whole cell lipidomic analysis, Δ-s-tether cells accumulate diacylglycerol (DAG) and exhibit reduced levels of phosphatidic acid (PA), phosphatidylserine (PS) as well as other phospholipids and sphingolipids. When the growth medium is supplemented with the lipid biosynthetic precursor choline, Δ-s-tether growth is comparable to WT, further indicating that defects in lipid metabolism are growth-limiting in cells lacking ER-PM contact. The importance of cER-PM MCSs for the regulation of lipid homeostasis is underscored by the finding that deletion of nonessential lipid regulatory genes, such as *OSH4* or *SAC1*, are lethal in Δ-s-tether cells, and that this lethality can be rescued by overexpression of phospholipid biosynthetic genes [15, 16]. Defects in cells lacking ER-PM tethers lead to constitutive activation of the inositol-attenuated Unfolded Protein Response (UPR) and the Environmental Stress Response (ESR) pathways, both of which are triggered by membrane damage [9, 15]. Although Δ-s-tether cells grow in the absence of cER-PM MCSs, their growth remains far from normal.

Phospholipid metabolism, specifically the ratio of DAG-to-PA levels within the nuclear ER membrane, is critical for regulating morphology and remodelling of the yeast nucleus prior to anaphase [17–27]. Unlike “open mitosis” in mammalian cells, nuclear segregation in budding yeast does not involve nuclear membrane breakdown. Instead, yeast chromosomes condense and attach to kinetochore microtubules within an intact nucleus [28, 29]. During the cell cycle, yeast nuclear morphology is dynamically regulated to maintain membrane integrity and facilitate nuclear segregation between mother cells and budding daughter cells during “closed mitosis” [25, 30, 31]. Despite obvious differences between open and closed mitosis, both processes share evolutionarily conserved mechanisms, underscoring the importance of phospholipid regulation in nuclear envelope segregation across eukaryotes [29, 32, 33].

In mammalian cells, controlled nuclear envelope breakdown occurs during open mitosis but nuclear ER membrane ruptures can be deleterious at other stages of the cell cycle when nuclei are intact. Such ruptures can contribute to human disease pathogenesis by causing chromosomal missegregation and genetic instability that promotes tumorigenesis [4, 5, 34, 35]. Even mechanical deformations of the intact nuclear membrane can disrupt cellular and nuclear functions [36, 37]. Nuclear envelope reformation and nuclear ER membrane ruptures are mended by a conserved mechanism involving the ESCRT-III (Endosomal Sorting Complex Required for Transport III) complex [38–41]. ESCRT-III is an oligomeric filament that binds to membranes to facilitate scission and repair [42, 43]. In this context, mechanisms for maintaining nuclear membrane integrity are conserved from yeast to humans, but the broader role of nuclear envelope integrity in preserving genomic stability remains unclear.

During budding yeast closed mitosis, metaphase expansion of the nuclear ER membrane adjacent to the nucleolus is orchestrated with subsequent actin-mediated “nucleopodial” projections of the nuclear membrane into the bud, after which cytoplasmic/astral microtubules pull the entire nucleus into the daughter bud during anaphase [44, 45]. Later in mitosis, alignment of the nucleus within the bud neck separating mother and daughter cells defines the axis for nuclear division during “karyokinesis” [46]. During nuclear growth and segregation, the Polo-like kinase Cdc5 links nuclear dynamics and cell cycle progression [47, 48]. Specifically, Cdc5 selects sites for nuclear ER membrane expansion and integrates Spindle Assembly Checkpoint (SAC) and the Spindle Position Checkpoint (SPOC) signaling with the mitotic exit network; therefore Cdc5 cell-cycle dependent translocations provide a readout for the proper coordination of nuclear processes with cell division in late mitosis [47–53].

Specific yeast tether proteins that establish cER-PM MCSs also facilitate the redistribution of cER during mitosis. The ER membrane proteins Scs2 and Ice2 are required for efficient inheritance of cER from the mother cell into the daughter bud, where cER is redistributed along the PM [54, 55]. Loss of both *SCS2* and *ICE2* causes significant defects in cER inheritance and disrupts septin organization at the bud neck, suggesting that Scs2 and Ice2 facilitate the regulated passage of cER across the diffusion barrier separating the mother cell from its daughter bud [55]. Despite the interconnection between cER and nuclear ER, however, deletion of *SCS2* and *ICE2* has no overt effect on nuclear ER [55]. Consistent with this fact, cER and nuclear ER are delivered to the daughter cell at different stages during the cell cycle and appear to rely on different mechanisms; notably, cER inheritance is microtubule independent [56]. Because of these distinct mechanisms of ER inheritance, the question arises whether cER association with the PM impacts either nuclear ER integrity or dynamics.

Here, we show that membrane tethers establishing yeast cER-PM MCSs collectively act as dual-function hubs that maintain nuclear envelope integrity by coordinating structural stabilization from intermembrane ER connections with phospholipid regulation within nuclear membranes. In cells lacking cER-PM MCSs, nuclear ER membrane integrity is severely compromised and cell survival is dependent on the nuclear ESCRT-III membrane repair complex. The lack of all cER-PM MCS tethers results in specific nuclear defects; chromosomal segregation remains unaffected but increased levels of spontaneous mutation suggest defects in genomic DNA repair. Mitotic nuclear dynamics are also impaired, leading to defects in nuclear segregation and an accumulation of cells with single undivided nuclei. These nuclear segregation defects trigger a SPOC-dependent cell-cycle delay, as evidenced by genetic interactions with SPOC regulators as well as Cdc5 spatiotemporal dysregulation. Together, these results highlight the structural importance of cER-PM attachment in maintaining nuclear membrane integrity and facilitating genomic segregation during mitosis.

## Results

### Nuclear ER membrane integrity is compromised in cells lacking cER-PM contact sites

Removing the membrane tethers that directly connect cER to the PM slows yeast growth and disrupts phospholipid synthetic flux [9, 14–16]. Given the importance of cER-PM MCSs in intermembrane signaling and lipid transfer, it is surprising that their elimination does not cause severe growth impairment. Although removal of cER-PM tethers is not lethal, it still causes significant cellular disruptions, particularly affecting ER morphology and the maintenance of the double ER membrane that forms the nuclear envelope (Fig 1).

**Figure 1.**
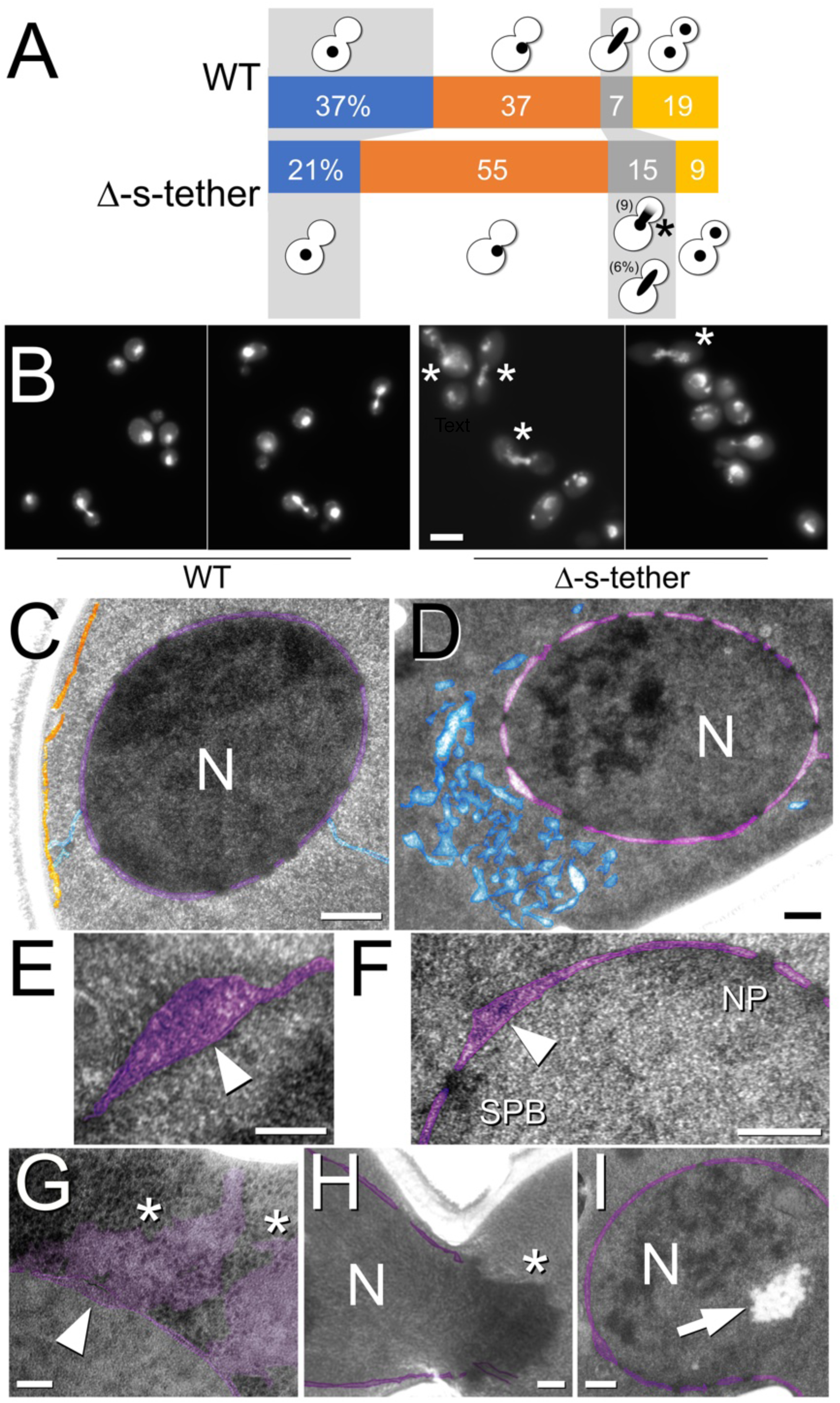
Defects in nuclear DNA and nuclear envelope structure in yeast cells lacking cER-PM MCSs. **(A)** Quantification of DAPI-stained nuclei in large-budded Δ-s-tether cells (CBY5838) (n = 163 cells) staged during the cell cycle according to nuclear morphology, relative to large-budded WT cells (SEY6210) (n = 80 cells). **(B)** Images of well-defined DAPI-stained nuclei in WT; diffusely fluorescent nuclei observed in Δ-s-tether cells during anaphase indicated by asterisks. Scale bar: 5 µm. **(C)** Transmission electron microscopy (TEM) of nuclei in WT, and **(D)** Δ-s-tether cells. Scale bars: 250 nm. **(E)** and **(F)** correspond to TEM images of nuclear membrane separations in Δ-s-tether cells. Arrowheads indicate membrane blisters forming between the inner and outer nuclear membranes. Scale bars: 100 and 250 nm, respectively. **(G)** TEM of erupted nuclear blisters in Δ-s-tether cells (arrowhead) accompanied by released material from the nuclear ER lumen, and potentially from within the nucleus (asterisks). Scale bar: 100 nm. **(H)** TEM of ruptured nucleus (asterisk) in anaphase Δ-s-tether cell. Scale bar: 250 nm. **(I)** TEM of amassed intranuclear lipid droplets (arrow). Scale bar: 250 nm. For TEM images, nuclear ER is false coloured in transparent purple; cytoplasmic ER in blue; cortical ER in orange. Nucleus, N; spindle-pole body, SPB; nuclear pore, NP.

The removal of cER-PM MCSs leads to a 2.6-fold increase in large-budded cells in exponential cultures, as compared to WT (n = 500 cells). When visualized by DAPI DNA fluorescence, a disproportionate number of these large-budded Δ-s-tether cells display an undivided nucleus lodged at or near the bud neck (Fig 1A and 1B). In contrast to WT cells where the nuclear DNA is clear and intact, half of Δ-s-tether cells exhibit a single anaphase nucleus with diffuse cytoplasmic DAPI fluorescence, suggesting DNA extrusion from the nucleus (Fig 1A and 1B). Upon closer inspection of cryofixed cells by transmission electron microscopy (TEM), Δ-s-tether cells exhibited several nuclear membrane defects not observed in WT cells (Fig 1C and 1D). In WT, the average width of the inner and outer nuclear membranes and the luminal nuclear periplasm between them is 26.7 ± 2.4 nm (n = 20 cells). In Δ-s-tether cells, however, the average distance expands to 34.2 ± 3.2 nm (n = 20 cells), consistent with observed increases in ER membrane separation induced upon UPR activation [57]. More extreme 50-90 nm separations of the inner and outer nuclear membranes are also apparent in 31% of Δ-s-tether nuclei (with one or more membrane “blisters” per sectioned nucleus), but not in WT nuclei (n > 31 cells) (Fig 1E and 1F). Although it is unclear whether these blisters always correspond to sites of nuclear envelope tearing, membrane ruptures at these sites are observed and are evocative of the DAPI-fluorescent DNA extrusions (Fig 1G). Although anomalous lipid droplet aggregates were observed in 34% of sectioned Δ-s-tether nuclei (n = 35 cells) (Fig 1I), similar to those reported in some phospholipid biosynthesis mutants [24], there was no correlation between the presence of intranuclear lipid droplets and membrane blisters/ruptures. Indeed, in Δ-s-tether cells, none of the nuclear blisters themselves contained nascent lipid droplets such as those seen within the perinuclear ER lumen in phospholipid mutants [24]. Notably, increased lipid saturation also results in significant nuclear envelope rupturing that correlates with an irregular distribution of nuclear pore complexes (NPCs) and spindle pole bodies (SPBs) [58]. Although these nuclear ER membrane defects mirrored those in cells lacking cER-PM tethers, parallel changes in NPC and SPB distributions were not observed. In addition to the defects observed in the nucleus and the nuclear envelope, the cytoplasmic ER in these cells also exhibits irregular membrane spacing and luminal expansion, relative to WT (Fig 1E and 1F). With respect to the inner and outer nuclear membranes, however, these findings indicated that the absence of ER-PM MCSs causes membrane structural distortions, which disrupt the organization and integrity of nuclear ER.

### ER-PM contact sites and ESCRT-III maintain nuclear integrity

To further investigate the severity of these nuclear ER membrane defects, we utilized benzyl alcohol, a membrane-fluidizing agent used in yeast to assess perturbations in nuclear ER membrane integrity and structure [59–61]. As a comparative control, the ESCRT-III mutant *snf7*Δ, which otherwise promotes membrane repair, is sensitive to 0.2% benzyl alcohol, whereas WT cells show no sensitivity [34, 38, 39, 43, 62, 63] (Fig 2A). To test if the removal of cER-PM MCSs induces similar nuclear ER membrane defects, Δ-s-tether cells were cultured in the presence of benzyl alcohol. As compared to WT, Δ-s-tether cell growth is significantly inhibited by benzyl alcohol similar to the *snf7*Δ control (Fig 2A). The hypersensitivity of Δ-s-tether cells to benzyl alcohol is consistent with the nuclear ER membrane deformations observed by TEM, suggesting nuclear envelope instability.

**Figure 2.**
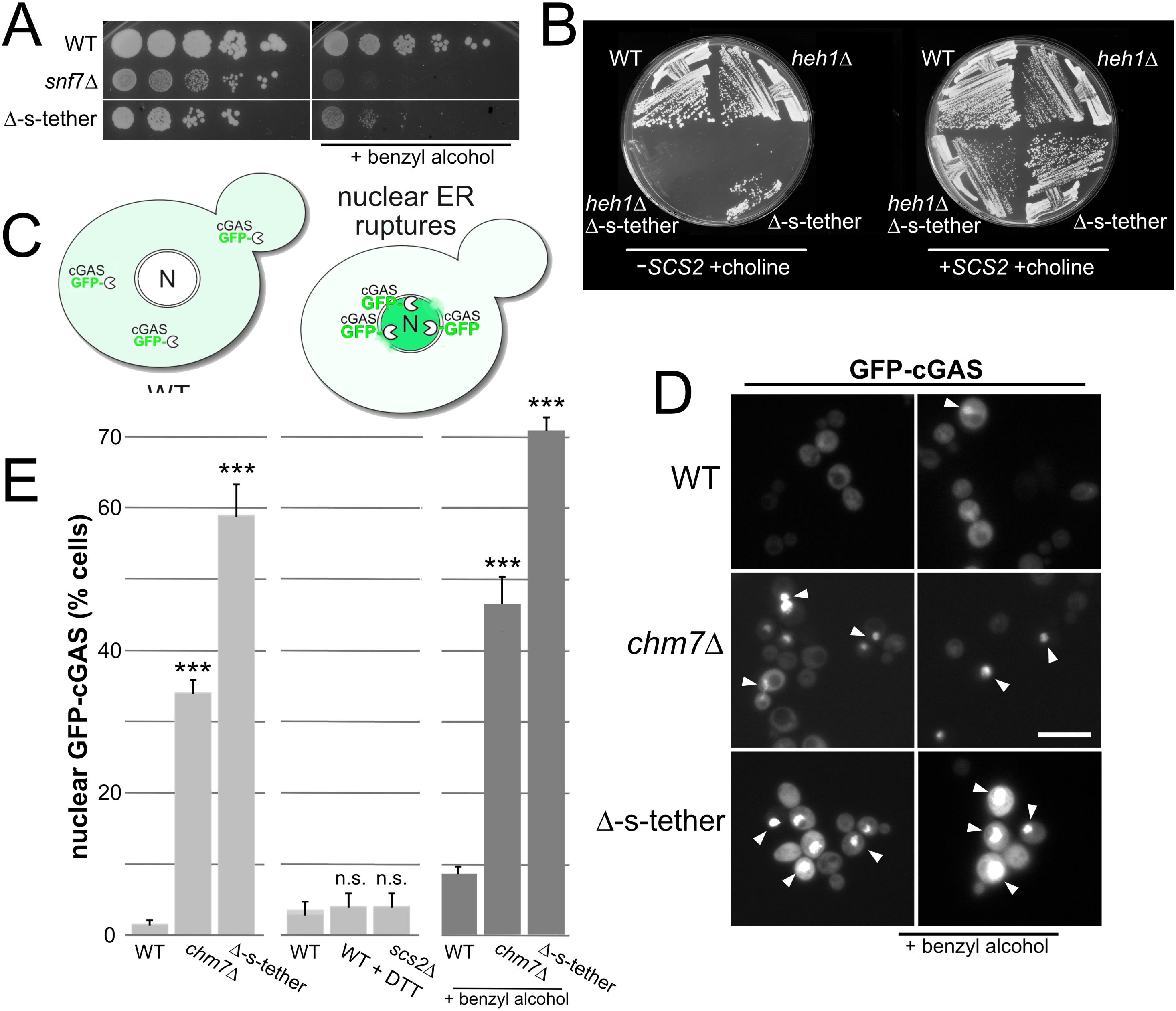
Nuclear membrane ruptures compromise nuclear integrity in cells without cER-PM tethers. **(A)** Sensitivity of Δ-s-tether cells to the nuclear membrane-destabilizing agent benzyl alcohol. Tenfold serial dilutions of WT (SEY6210), Δ-s-tether (CBY5838), and *snf7*Δ (CBY7704) cultures spotted onto solid rich medium with or without 0.2% benzyl alcohol and incubated for 4 days at 30°C. **(B)** WT (SEY6210), Δ-s-tether (CBY5838), *heh1*Δ (CBY8699), and *heh1*Δ Δ-s-tether (CBY8701) cells, with selection for (+*SCS2*) or against (-*SCS2*) a complementing plasmid (pSCS2), streaked onto solid growth medium supplemented with and without 1 mM choline incubated for 4 days at 30°C. **(C)** GFP-cGAS fluorescence assay to measure nuclear envelope ruptures. GFP-cGAS fluorescence is normally diffuse through the cytoplasm but exposure of nuclear double-stranded DNA to otherwise cytoplasmic GFP-cGAS results in bright fluorescence within nuclei and sites at the nuclear membrane periphery. **(D)** Representative images of GFP-cGAS (pCB1465) expressed in WT, *chm7*Δ (CBY8697), and Δ-s-tether cells cultured at 30°C with or without 0.5% benzyl alcohol (as indicated) showing cytoplasmic versus nuclear localization corresponding to ruptured nuclear membranes. Arrowheads indicate examples of nuclear GFP-cGAS accumulation. Scale bar: 10 µm. **(E)** Bar graphs quantifying nuclear GFP-cGAS fluorescence in WT, *chm7*Δ, and Δ-s-tether cells cultured at 30°C with or without benzyl alcohol. The lack of nuclear GFP-cGAS fluorescence in *scs2*Δ (CBY9065) cells and in WT cells treated within 2 mM DTT for 4 hr at 30°C is also quantified. n.s. = not statistically significant relative to WT; \*\*\**p* ≤ 0.004 (n ≥ 200 cells/strain).

We hypothesized that, in the absence of cER-PM MCSs, cell growth persists because ESCRT-III repairs nuclear membrane ruptures that would, presumably, be lethal. Indeed, the deletion of *SNF7* (or deletion of other ESCRT-III mutants) has been reported to exacerbate growth defects of yeast lacking a subset of cER-PM tethers (64); consistent with this finding, we were unable generate viable *snf7*Δ Δ-s-tether cells. Furthermore, the deletion of *HEH1*/*SRC1* in Δ-s-tether cells is also lethal (Fig 2B) (S1 Table). Heh1 is an adaptor that localizes and recruits ESCRT-III to the inner nuclear membrane surface, where it repairs membrane damage and safeguards nuclear envelope integrity [65, 66]. These findings implicate both cER-PM MCSs and Heh1-ESCRT-III in the maintenance of nuclear membrane stability.

To quantify how cER-PM MCSs specifically impact yeast nuclear integrity, we assayed nuclear ER membrane ruptures by analyzing the nuclear enrichment of a GFP-cyclic GMP-AMP synthase fusion protein (GFP-cGAS) (Fig 2C). In mammalian cells, the cGAS-STING pathway detects foreign cytoplasmic DNAs and responds by inducing innate immune responses [67, 68]. When expressed in yeast, we used the binding of GFP-cGAS to double-stranded DNA to detect when cytoplasmic GFP-cGAS penetrates nuclei through nuclear membrane ruptures. In WT, GFP-cGAS fluorescence is mainly observed as a cytoplasmic background fluorescence, but a small increase in nuclear GFP-cGAS is detected after the addition of 0.2% benzyl alcohol to the culture medium for 1 hr (Fig 2D and 2E). In humans, ESCRT-III is recruited by CHMP7 to repair nuclear envelope ruptures, and its yeast homologue is encoded by *CHM7* [34, 39, 43, 62, 63, 69–71]. As a control for detecting nuclear ruptures, we deleted *CHM7* and analyzed its impact on nuclear GFP-cGAS accumulation with and without benzyl alcohol. GFP-cGAS fluorescence concentrates within nuclei in about a third of *chm7*Δ cells, with a further nuclear accumulation upon benzyl alcohol addition (Fig 2D and 2E). In comparison, a much greater percentage of Δ-s-tether cells show intense nuclear GFP-cGAS fluorescence, which also increases with benzyl alcohol addition (Fig 2D and 2E). Even though Scs2 represents the tether that contributes most to cER attachment with the PM [9], the deletion of *SCS2* by itself had no significant effect on nuclear membrane integrity as detected by nuclear GFP-cGAS (Fig 2E). Moreover, while chronic ER stress and UPR induction has been presumed to cause nuclear ER ruptures, and removing multiple cER-PM tethers triggers the UPR pathway [9, 14, 15], no significant effect on the nuclear accumulation of GFP-cGAS was detected upon extended treatment of WT cells with 2 mM DTT for 4 hours (Fig 2E); far exceeding the time required for complete UPR induction [72]. These results indicate that collectively all cER-PM tethers are needed to ensure nuclear membrane stability and nuclear membrane rupturing is not an inherent consequence of ER stress. To determine when during the cell cycle nuclear rupturing occurs in Δ-s-tether cells, we used 4-D fluorescence microscopy to track when nuclear GFP-cGAS tears appear in mitotically growing cells (S1 Fig) (S1 Video). Because spontaneous nuclear GFP-cGAS could be visualized throughout the cell cycle, nuclear rupturing in Δ-s-tether cells does not appear to be exclusively a consequence of physical stresses exerted on anaphase nuclei when the nuclear envelope is pulled between mother and daughter cells.

To directly visualize Chm7-ESCRT-III recruitment to areas of nuclear membrane tears, Chm7-mRuby2 and GFP-cGAS were expressed together in WT and Δ-s-tether cells. In general, nuclear GFP-cGAS and Chm7-mRuby2 foci are rarely detected in WT cells (Fig 3A and 3B). In Δ-s-tether cells, however, almost half of cells with nuclear GFP-cGAS have overlapping Chm7-mRuby2 fluorescence (Fig 3A and 3B). In the absence of cER-PM MCSs, the observed colocalization of Chm7 with GFP-cGAS, though not complete, suggests that nuclear Chm7-ESCRT-III facilitates membrane repair.

**Figure 3.**
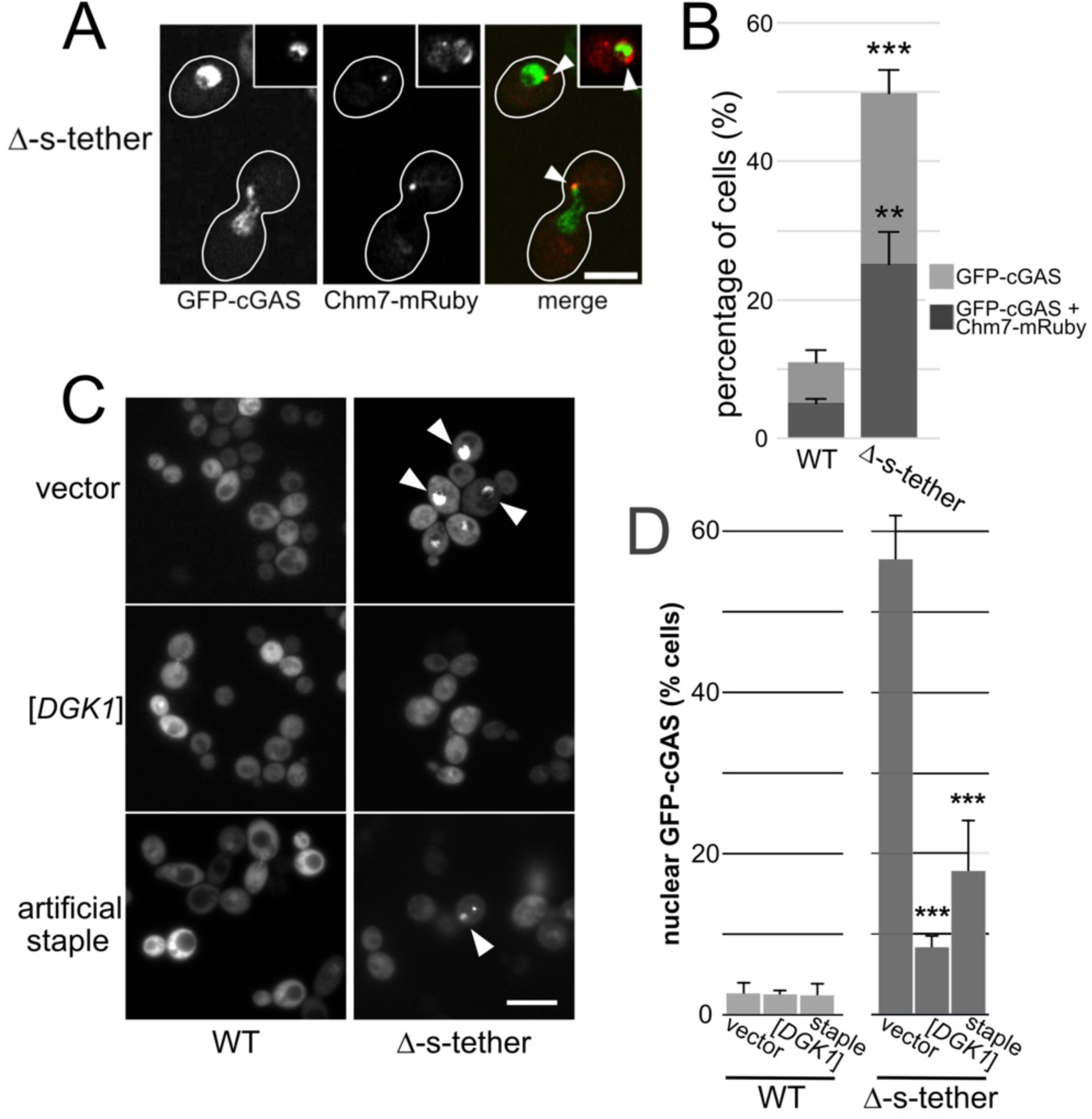
Nuclear ESCRT-III repairs nuclear envelope ruptures in cells lacking cER-PM MCSs. **(A)** Representative confocal fluorescence microscopy images of GFP-cGAS (pCB1465) nuclear membrane ruptures colocalized with Chm7-mRuby (pCB1661) coexpressed in Δ-s-tether (CBY5838) cells at 30°C. Cell peripheries are traced as shown, and the insert shows an anaphase GFP-cGAS fluorescent nucleus with accumulated Chm7-mRuby along the nuclear membrane. Scale bar: 5 µm. **(B)** Bar graph quantifying relative percentages of Chm7-mRuby colocalization with GFP-cGAS in WT (SEY6210) or Δ-s-tether cells (n ≥ 300 cells/strain), using wide-field fluorescence imaging. \*\**p* = 0.006;\*\*\**p* < 3 × 10^-5^ (as compared to the Δ-s-tether vector control). **(C)** Suppressors of Δ-s-tether growth defects also rescue nuclear rupturing shown by GFP-cGAS assay. Representative images using wide-field fluorescence microscopy of GFP-cGAS expressed in WT and Δ-s-tether cells transformed either with a vector control (YEplac181), multicopy *DGK1* (pCB1346) or artificial staple (pCB1188) plasmids. **(D)** Bar graph quantifying relative percentages of GFP-cGAS-fluorescent ruptured nuclear membranes corresponding to images in C (n ≥ 600 cells/strain). \*\*\**p* ≤ 0.003 as compared to the Δ-s-tether vector control.

### Parallel mechanisms rescue nuclear membrane integrity defects in the absence of ER-PM MCSs

The deletion of specific lipid regulatory genes (e.g. *OSH4* and *SAC1*) is lethal in Δ-s-tether cells [14–16]. This lethality can be suppressed by two independent mechanisms: restoring cER-PM contact through expression of a non-specific “artificial cER-PM membrane staple,” or enhancing phospholipid synthetic flux [14–16]. Artificial staple expression in Δ-s-tether cells re-establishes cER association with the PM, which rescues growth defects and significantly suppresses nuclear rupturing, as shown by decreased nuclear GFP-cGAS foci (Fig 3C and 3D) [14]. By lipidomic analysis, it has been shown that the artificial staple has little impact on the altered phospholipid composition in Δ-s-tether cells [15]. In contrast, increased expression of the Dgk1 diacylglycerol kinase, which synthesizes the phospholipid precursor PA from DAG, does not restore direct cER-PM association in cells lacking cER-PM tethers. However, multicopy *DGK1* expression does restore phospholipid flux and Δ-s-tether cell growth [14, 16]. As shown by GFP-cGAS assay, nuclear membrane ruptures in Δ-s-tether cells, are also restored by *DGK1* overexpression (Fig 3C and 3D) [14–16]. Through distinct mechanisms, nuclear ER structure and integrity are supported by both balanced membrane composition and non-specific physical stabilization by cER-PM tethering, that might cradle the nucleus against mechanical stresses.

Human “nuclear envelopathies” are chiefly represented by mutations affecting nuclear structural support (e.g. nuclear lamins), but also by those associated with lipodystrophies (e.g. BSCL2 [seipin] and AGPAT2) that affect DAG and PA acid synthesis [73]. In *S. cerevisiae*, the balance between DAG and PA is an important determinant of nuclear membrane morphology, where DAG synthesis by the lipin homologues Pah1 is in opposition to PA synthesis by Dgk1 [20, 21, 74]. The DAG:PA balance within the inner nuclear membrane influences nuclear envelope outgrowth by regulating phospholipid synthesis adjacent to the nucleolus, which is required for nuclear membrane expansion prior to cell and nuclear division [19, 33, 75]. Because *DGK1* overexpression rescues Δ-s-tether growth defects, we predicted the opposite, namely *PAH1* deletion, would also suppress. Not only does *pah1*Δ restore normal growth to Δ-s-tether cells, but deletion of *NEM1*, which encodes a phosphatase regulatory subunit that activates Pah1p, also rescued growth defects (Fig 4A) (Table1). Given that *DGK1* overexpression and *pah1*Δ suppress Δ-s-tether growth defects, and Δ-s-tether cells exhibit reduced PA and elevated DAG levels, this suggested that PA synthesis is inhibited in Δ-s-tether cells. However, whole cell imaging using genetically encoded fluorescent PA (GFP-Spo20^51-91^) and DAG (C1δ-GFP) probes revealed little change in their distributions in Δ-s-tether cells: PA is enriched at the cell cortex, while DAG localization is mainly in the vacuole and PM [16]. To specifically examine nuclear PA and DAG in WT and Δ-s-tether cells, the lipid-binding probes were targeted to the inner nuclear membrane via added nuclear localization sequences (NLSs) (Fig 4B) [24]. To our surprise, in Δ-s-tether cells the fluorescence of the nuclear-targeted PA-mCherry probe (NLS-Opi1 Q2-mCherry) increased by an average of 2.5-fold, relative to WT (Fig 4B and 4C). In contrast, the NLS-DAG-GFP probe (NLS-C1a+C1b-GFP) remains at the PM and vacuole and is only partially distributed to the nuclear envelope in WT cells, but nuclear membrane localization is more evident in Δ-s-tether cells. Relative to WT cells, in Δ-s-tether cells NLS-DAG-GFP nuclear envelope fluorescence is 4.7-fold more intense, indicating DAG accumulation within the inner nuclear membrane (Fig 4B and 4C). Although the relative nuclear fluorescence of both probes increased in Δ-s-tether cells, the PA:DAG ratio is still half of that observed in WT cells, indicating dysregulation of the PA and DAG nuclear membrane balance. Perhaps even more notable, however, is the appearance of highly motile intranuclear DAG-enriched particles that likely correspond to the nuclear lipid aggregates observed by TEM in Δ-s-tether cells (Fig 1I) (S2 Video). Using the NLS-PA probe to confirm nuclear localization, these intranuclear DAG particles were seen in 81% of nuclei in Δ-s-tether cells and in only 2.6% of WT cells (n = 90 cells per strain). Despite the potential for violent collisions between these particles and inner nuclear membrane there is, however, no evidence that they compromise nuclear envelope integrity. In Δ-s-tether cells, *DGK1* overexpression restores phospholipid flux and corrects the nuclear NLS-PA-mCherry intensity to WT levels, whereas the artificial staple has little effect, and NLS-PA-mCherry fluorescence remains significantly higher (Fig 4D and 4E). Consistent with previous lipidomic analysis of Δ-s-tether cells expressing the artificial staple, non-specific reattachment of cER to the PM does not normalize PA levels or nuclear distribution [14, 16]. These results provide additional evidence that cER-PM tethering mediates two independent mechanisms that together contribute to nuclear ER membrane stability.

**Figure 4.**
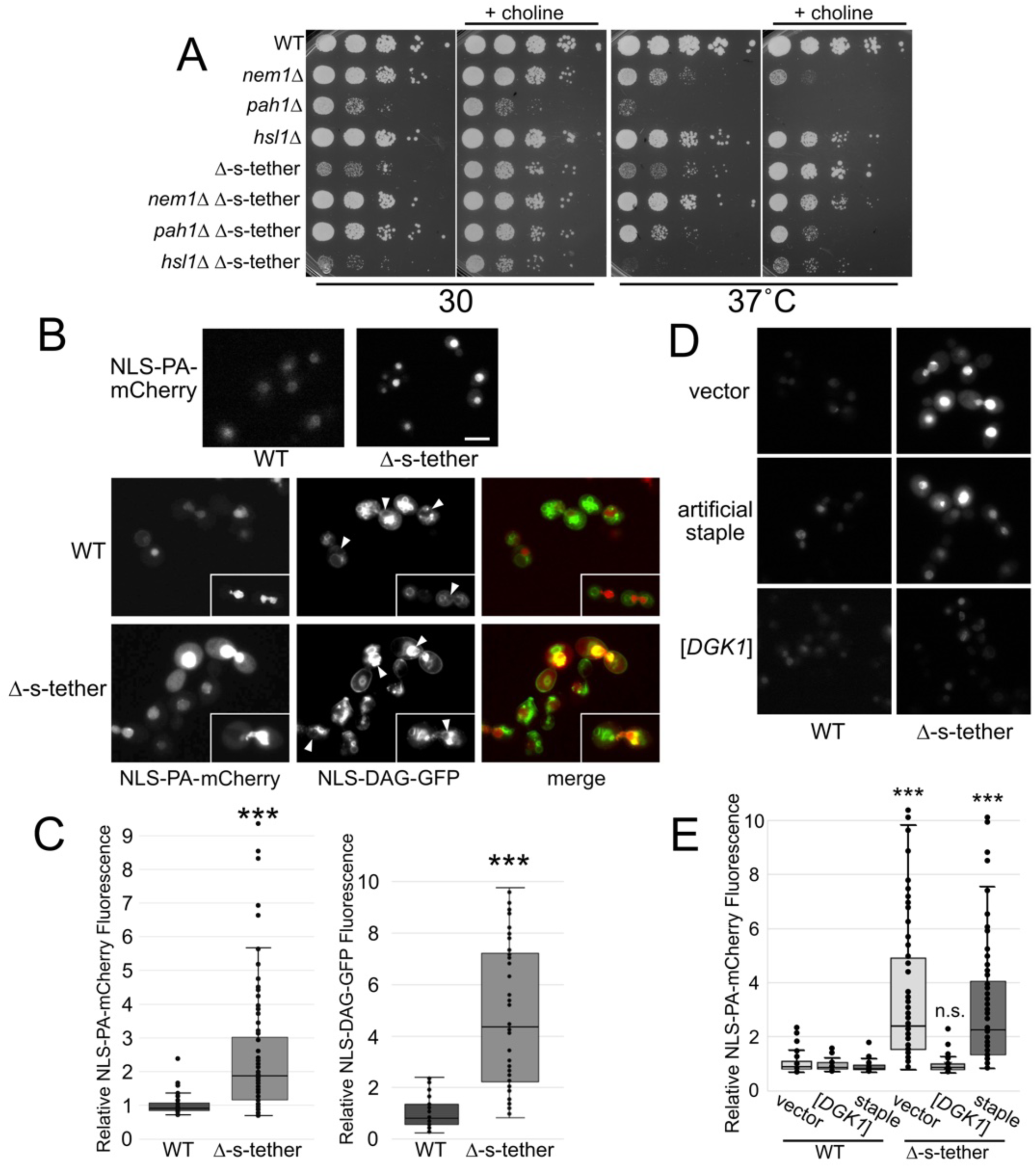
Nuclear PA and DAG increase when cER-PM tethering is eliminated. **(A)** Tenfold serial dilutions of WT (SEY6210), *nem1*Δ (CBY8705), *pah1*Δ (CBY8707), *hsl1*Δ (CBY8731), Δ-s-tether (CBY5838), *nem1*Δ Δ-s- tether (CBY8709), *pah1*Δ Δ-s-tether (CBY8711), and *hsl1*Δ Δ-s-tether (CBY8739) cultures spotted onto solid growth medium incubated for 3 days at 30°C. **(B)** Top: Representative images of WT and Δ-s-tether cells expressing the fluorescent nuclear PA probe (pRS316-NLS-Opi1-Q2-mCherry). Bottom: the same cells coexpressing the nuclear PA probe and the nuclear DAG probe (pRS316-NLS-C1a+C1b-GFP). Cells were cultured in liquid selective growth medium at 30°C. Scale bar: 10 µm. **(C)** Box and whiskers plots showing quantification and range of fluorescence corresponding to the coexpression of the nuclear NLS-PA-mCherry probe (n ≥ 50 cells) or NLS-DAG-GFP probe (n = 50 cells) in WT and Δ-s-tether cells. Fluorescence values are normalized to the mean WT intensity. \*\*\**p* ≤ 1.2 × 10^-12^. **(D)** Representative images using wide-field fluorescence microscopy of NLS-PA-mCherry expressed in WT and Δ-s-tether cells transformed either with a vector control (YEplac181), multicopy *DGK1* (pCB1346) or artificial staple (pCB1188) plasmids. **(E)** Box and whiskers plots quantifying relative proportions of NLS-PA-mCherry fluorescence (normalized to mean WT intensity) corresponding to images in D (n ≥ 200 cells). High-copy *DGK1* rescues nuclear NLS-PA-mCherry accumulation in Δ-s-tether nuclei, but the artificial staple does not. n.s. = not statistically significant; \*\*\**p* ≤ 0.001 as compared to the Δ-s-tether vector control.

### Elimination of cER-PM MCSs leads to genomic instability and disrupts mitotic nuclear segregation

Nuclear ER ruptures disrupt cell division and genomic stability, which can lead to heritable genetic change [37, 76]. Nuclear rupture in mammalian cells, arising from either lamin defects or morphological stress on nuclei during confined cellular migration, can result in chromothripsis-like chromosome rearrangements and DNA damage [4, 5, 34, 73, 77]. Indeed, even minor nuclear membrane deformations in cancer cells seem sufficient to elicit replication stress and DNA damage [78]. Nuclear envelope rupture in cancer cells has also been proposed to drive mutagenesis through the efflux of DNA repair proteins from nuclei and/or influx of potentially damaging cytoplasmic proteins [79–81]. In this context, as shown by the GFP-cGAS assay, cytoplasmic protein influx into the nucleus is readily detected in most Δ-s-tether cells. To determine if the absence of ER-PM MCSs contributes to chromosome instability, a centromeric (*CEN*) plasmid loss assay was used to test for chromosome loss in Δ-s-tether cells. In contrast to the *chl4*Δ control, which undergoes chromosome mis-segregation due to defective kinetochore attachment [82], the nuclear ER membrane defects in Δ-s-tether cells do not contribute to *CEN* plasmid loss, indicating an absence of chromosomal instability (Fig 5A). Beyond chromosome segregation, nuclear envelope integrity and compartmentalization affect mutagenic frequency, as demonstrated by the hypersensitivity of ESCRT-III mutants to phleomycin-induced DNA damage [83]. To investigate whether nuclear membrane ruptures cause defective DNA repair and increase mutagenic frequency, fluctuation tests were conducted to determine relative spontaneous mutation rates using a plasmid read-out system [84]. Relative to WT cells, Δ-s-tether cells transformed with a *CEN* plasmid containing both *URA3* and *LEU2* exhibited a significantly higher frequency of spontaneous *URA3* mutations that confer resistance to 5-fluoroorotic acid (5-FOA) in transformants that still retain *LEU2*-dependent leucine prototrophy (Fig 5B). As predicted, the *rad52* mutator strain control generated even higher levels of spontaneous mutations (Fig 5B).

**Figure 5.**
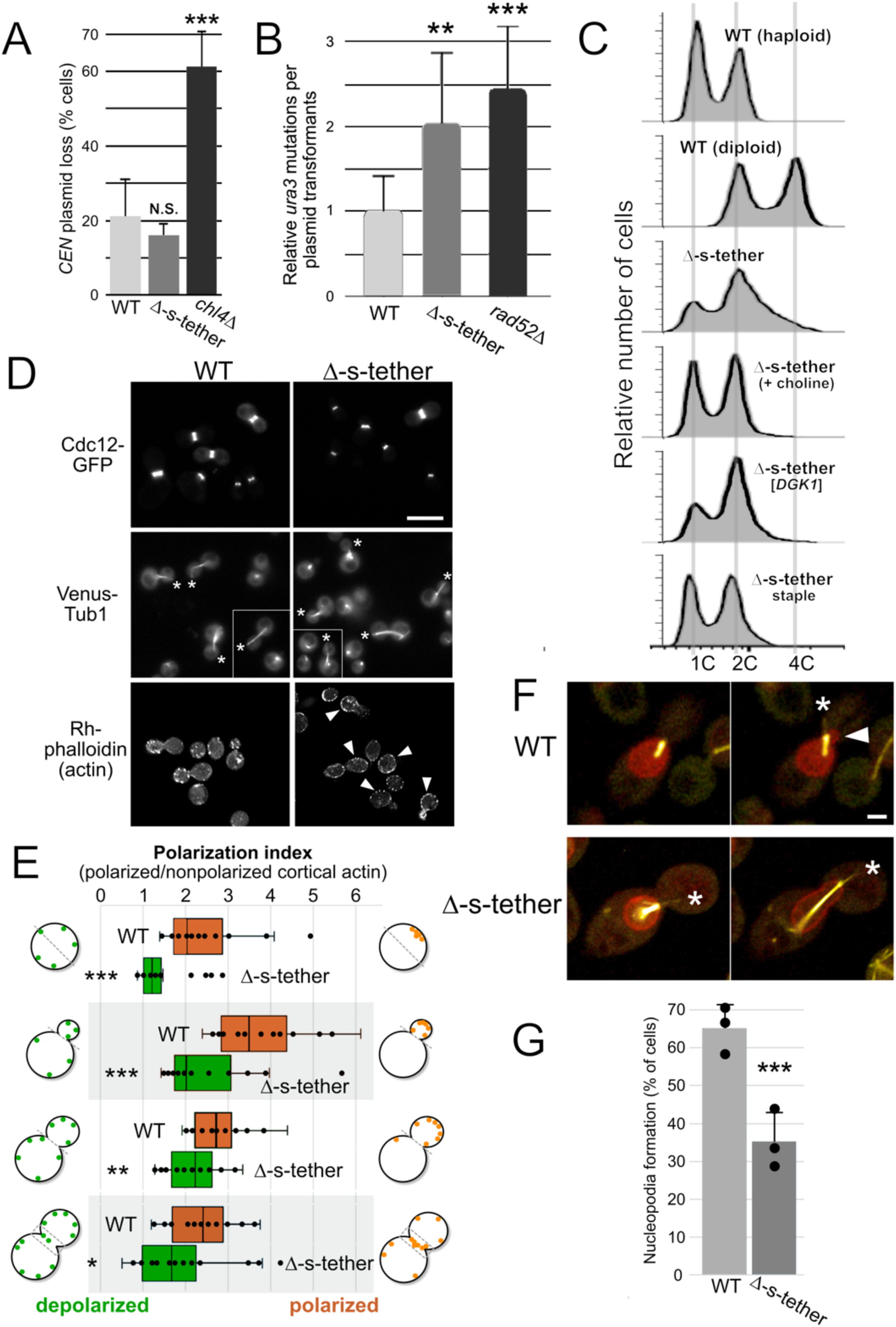
Elimination of cER-PM MCSs interferes with genomic DNA repair and mitotic nuclear dynamics. **(A)** Centromere (*CEN*) plasmid loss assay. Relative percentages of *CEN* plasmid (pRS416) retention in WT (SEY6210), Δ-s-tether (CBY5838), and *chl4*Δ (CBY8713) after overnight growth in non-selective liquid medium at 30°C and spread onto selective versus non-selective solid media for 4 days at 30°C. **(B)** Fluctuation tests quantifying the relative level of spontaneous 5-FOA-resistant *ura3* mutations per *URA3*/*LEU2*/*CEN* plasmid (pCB1774) transformants in WT, Δ-s-tether, and *rad52*Δ (CBY8597) cells. Mutation rates were calculated using the Ma–Sandri–Sarkar maximum likelihood (MSS-MLE) method, error bars represent 95% confidence intervals (n = 89 independent cultures for each strain). \*\**p* = 0.007; \*\*\**p* = 1 × 10^-4^ (as compared to WT). **(C)** Analysis of genomic inheritance by flow cytometry: DNA content as measured by propidium iodide fluorescence of WT haploid (SEY6210), WT diploid (CBY57), and Δ-s-tether (CBY5838) cells transformed with the vector control (YEplac181). Also shown are Δ-s-tether cells transformed with either the high-copy *DGK1* suppressor (pCB1346) or the artificial staple (pCB1185). Asynchronous log-phase cultures were cultured at 30°C in the absence of choline unless otherwise indicated (+choline). Relative to the area of propidium fluorescence, ploidy (1-4C) of cell populations is as shown. **(D)** Representative images of WT and Δ-s-tether cells grown at 30°C expressing the septin Cdc12-GFP (pCB502), or Venus-Tub1 (CBY6095 and CBY6107 integrant strains) to show astral and intranuclear microtubules, or stained with Rhodamine-phalloidin to visualize cortical actin patches. Asterisks denote proper polarization of astral microtubules into the bud; arrowheads indicate actin patch depolarization spreading into the mother cell; Rhodamine-phalloidin actin patches in Δ-s-tether cells are smaller, less fluorescent, and more numerous. Scale bar: 10 µm. **(E)** Box and whiskers plots showing quantification of cortical actin patch polarization in WT and Δ-s-tether at different cell-cycle stages (unbudded, small-, medium-, and large-budded). The polarization index is defined as the ratio of fluorescence localized at areas of polarization (indicated by hatched lines in the cell schematics) to the remaining cortical fluorescence (n = 20 cells per stage per strain). \*\*\**p* < 0.0005, \*\**p* = 0.008, \**p* = 0.1. **(G)** Representative images of asynchronous log-phase WT (CBY6095) and Δ-s-tether (CBY6107) cells grown at 30°C expressing Venus-Tub1 and transformed with a plasmid expressing the nuclear NLS-PA-mCherry probe (pRS316-NLS-Opi1-Q2-mCherry). Images show first and last frame of videos tracking nuclear membrane outgrowth (red) into the bud relative to microtubules (yellow). WT cells exhibit nucleopodia shortly before nuclear migration into the bud, whereas nucleopodia are infrequently observed in Δ-s-tether cells, even upon observing them until the onset of anaphase (as shown). The arrowhead indicates microtubule-independent nucleopodia; asterisks denote the direction of microtubule extension into the bud. Scale bar: 2 µm. **(H)** Bar graph quantifying nucleopodia formation in WT and Δ-s-tether cells coexpressing Venus-Tub1 and the nuclear NLS-PA-mCherry probe. Nucleopodial outgrowths were scored when the nuclear membrane expanded into the bud, independent of the direction of polarized microtubule extension (grand mean calculated from n = 5 cell videos for each of 3 independent trials). Note that only Δ-s-tether cells that overcome the cell-cycle delay and proceed into anaphase were scored. \*\*\**p* = 0.007

Taken together, these results suggest that eliminating cER-PM MCSs does not increase chromosomal instability but instead interferes with the repair of spontaneous mutations (or directly induces genomic mutation).

To determine whether mitotic nuclear segregation is disrupted in Δ-s-tether cells, we analyzed nuclear DNA content in asynchronous log-phase cultures using flow cytometry. Consistent with the increased proportion of large-budded cells indicative of a cell-cycle delay prior to cell division, Δ-s-tether cells accumulated multiploid (C ≥ 2) cells with a corresponding reduction in cells with a G1 haploid (C = 1) DNA content (Fig 5C). Suppressors of Δ-s-tether growth defects restored normal haploid DNA profiles, especially choline supplementation and the artificial staple, though high-copy *DGK1* expression had a more moderate effect (Fig 5C). Despite increased genome equivalents, preanaphase Δ-s-tether cells predominantly contain a single DAPI-stained nuclear mass; binucleate nuclei are observed only in a small subset of cells and almost never seen in WT (Fig 1A and S3 Fig). Multiploid nuclei contained within a single nuclear envelope suggest a defect in progression through mitotic nuclear division (karyokinesis).

Cell-cycle dependent changes in nuclear segregation and dynamics are often driven by changes in cytoskeletal assembly and organization [85]. In budding yeast, the polarized positioning of astral microtubules directs nuclear migration towards the bud, while actin-dependent nucleopodia initiate nuclear ER membrane outgrowth from the mother cell through the bud neck into the bud [44, 56, 86]. At the bud neck, septin filaments position the actomyosin ring for PM constriction to coordinate cytokinesis with nuclear division [87]. In Δ-s-tether cells, distributions of Venus-Tub1 and Cdc12-GFP fluorescence indicate that microtubule organization and polarization, and bud-neck septin filament assembly, remain similar to WT (Fig 5E). The normal distribution of Cdc12-GFP was surprising given that the combined deletions of *SCS2* and *ICE2* cause septin mislocalization [55]. This might suggest that the other tethers counterbalance Scs2 and Ice2 effects on septins. In contrast, actin patches, which are normally concentrated within the daughter bud, exhibit either partial or significant depolarization within Δ-s-tether cells, depending on cell-cycle morphological stage (Fig 5D and 5E). Because nucleopodial extensions into the bud are actin-dependent, these findings led us to hypothesize that actin patch depolarization in Δ-s-tether cells disrupts nucleopodial protrusions. This is also consistent with the accumulation of large-budded Δ-s-tether cells, many of which contain undivided anaphase nuclei lodged near the bud neck that fail to translocate into daughter cell, despite having no detectable defect in microtubule polarization.

To directly visualize nucleopodial dynamics, we analyzed preanaphase cells co-expressing Venus-Tub1 and NLS-PA-mCherry to monitor the direction of microtubule extension relative to nuclear envelope outgrowth into the daughter bud. Although nucleopodia are most readily observed in cells after release from a G2/M cell-cycle arrest [44], we sought to avoid complications with additional cell-cycle manipulations. Therefore, we tracked microtubule-independent nuclear membrane outgrowth into buds in untreated, asynchronously cultured WT and Δ-s-tether cells using 4D super-resolution fluorescence microscopy.

In our analysis, almost all detectable nucleopodia form within 12 min after the nucleus reaches the bud neck in pre-anaphase cells. By tracking cells during this period, we found that microtubule-independent nuclear outgrowth occurs in 95% of all WT cells (n = 20 cells). In contrast, nucleopodia formation was significantly reduced in large-budded Δ-s-tether cells whose nuclei remained lodged at the bud neck throughout the 12 min observation period, occurring only in only 25% of cells (n = 20 cells; *p* = 5 × 10⁻⁶). Even in those Δ-s-tether cells where nuclei did traverse the bud neck, microtubule-independent nucleopodial outgrowths into the bud remained significantly less frequent than in WT (S3 Video) (Fig 5F and 5G). In the absence of cER-PM MCSs, impaired cortical actin patches polarization and/or dysregulation of nuclear membrane phospholipids, both of which promote nucleopodial membrane outgrowth [44], likely contribute to the observed defects in nuclear segregation and karyokinesis.

### Defects in cER-PM MCSs trigger a Spindle Position Checkpoint (SPOC) cell-cycle delay

Defects in nuclear segregation can activate checkpoint mechanisms that pause or arrest the cell cycle, which allows time for genome repair before proceeding [51, 88]. Because yeast lacking cER-PM MCSs accumulate large-budded cells, we hypothesized that a SAC and/or SPOC cell-cycle delay might be induced to provide additional time for proper nuclear positioning or ER membrane resealing to avoid potentially catastrophic consequences later during mitotic exit. Although Δ-s-tether cells show no defect in the *CEN* plasmid loss assay defect, suggesting that SAC checkpoint monitoring of chromosome segregation is unaffected, we tested if disruption of SAC affects growth in cells lacking cER-PM MCSs. Although deletion of the SAC regulator *BUB3* confers sensitivity to growth at low temperatures and the microtubule inhibitor benomyl, both of which inhibit spindle formation [89], *BUB3* deletion in Δ-s-tether cells does not exacerbate this sensitivity (Fig 6A). Indeed, in the presence of both benomyl and lower temperature, *bub3*Δ is no longer lethal when combined with Δ-s-tether mutations (Fig 6A). Furthermore, the deletion of *CHL4,* which elicits a SAC delay, has a negligible effect when deleted in Δ-s-tether cells (Fig 6A) (S1 Table). Because the deletion of SAC regulators has little effect on Δ-s-tether cells, these findings suggest that loss of ER-PM contacts does not trigger a SAC-dependent cell-cycle delay. In contrast, *BUB2* deletion impairs the SPOC response and causes synergistic growth defects in cells lacking cER-PM MCSs when cultured at low temperature in the presence of benomyl (Fig 6A). Moreover, in Δ-s-tether cells, deletions of several other SPOC regulators, including multifunctional kinases that also regulate phospholipid regulation, lead to severe growth defects or lethality (S1 Table). Because microtubule polarization remains unaffected in Δ-s-tether cells (Fig 5E), these findings suggest that SPOC also monitors other microtubule-independent nuclear events including those impacted by cER-PM tethering.

**Figure 6.**
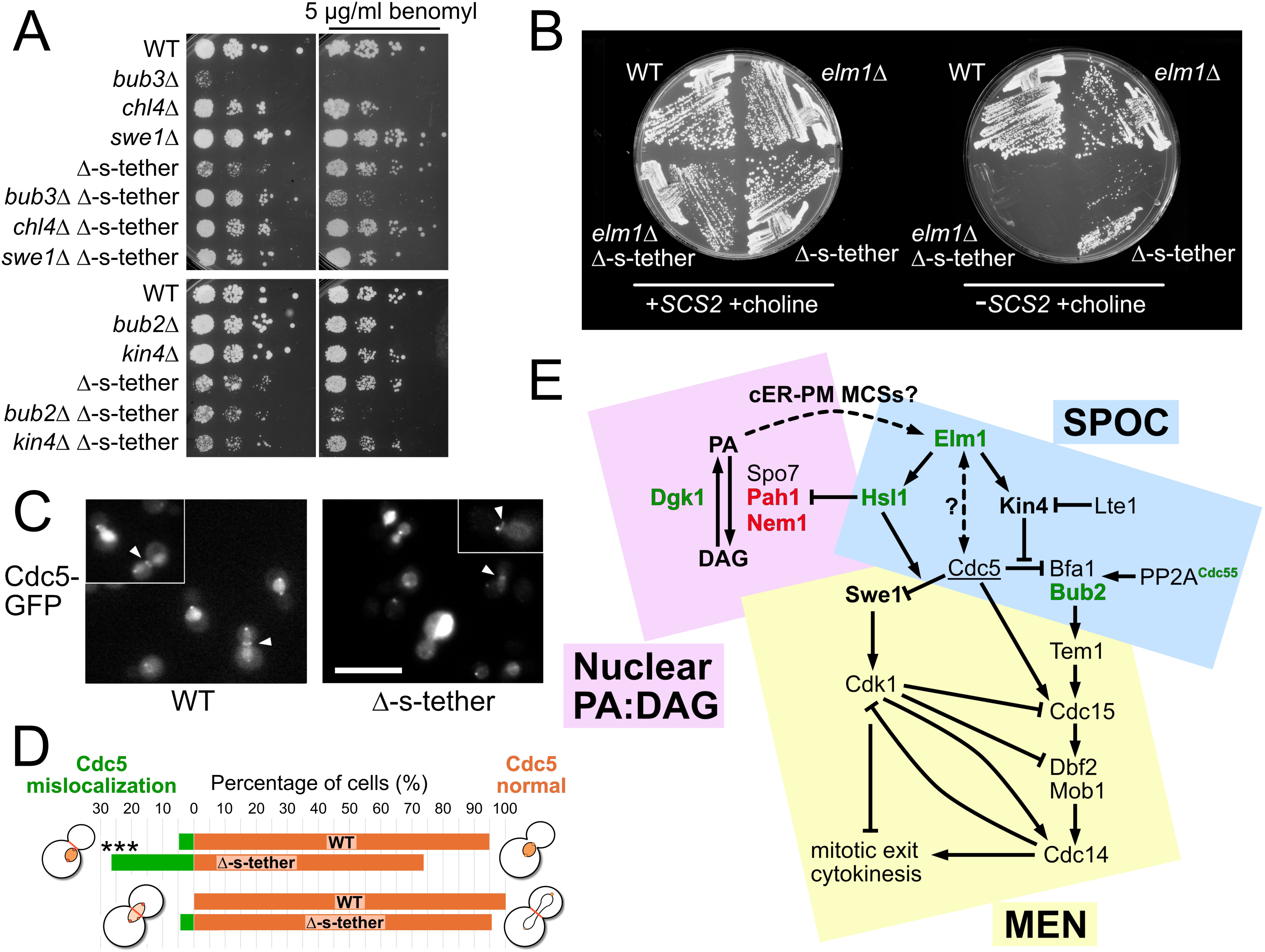
The Spindle Position Checkpoint (SPOC) monitors the nuclear membrane in response to dysfunctional cER-PM tethering. **(A)** Tenfold serial dilutions (top) of WT (SEY6210), *bub3*Δ (CBY8715), *chl4*Δ (CBY8713), *swe1*Δ (CBY8733), Δ-s-tether (CBY5838), *bub3*Δ Δ-s-tether (CBY8717), *chl4*Δ Δ-s-tether (CBY8719), *swe1*Δ Δ-s-tether (CBY8741) cells showing the comparative lack of growth defects between Δ-s-tether and SAC- associated mutations; and serial dilutions (bottom) of WT, *bub2*Δ (CBY8721), *kin4*Δ (CBY8723), Δ-s-tether, *bub2*Δ Δ-s-tether (CBY8725), *kin4*Δ Δ-s-tether (CBY8727) cells indicating SPOC-associated synthetic growth defects. All cultures were spotted onto solid growth medium incubated for 14 days at 16°C in the presence and absence of 5 µg/mL benomyl. **(B)** WT (SEY6210), Δ-s-tether (CBY5838), *elm1*Δ (CBY8729), and *elm1*Δ Δ-s-tether (CBY8737) cells, with selection for (+SCS2) or against (-SCS2) a rescuing plasmid (pSCS2), streaked onto solid growth medium supplemented with 1 mM choline incubated for 4 days at 30°C. **(C)** Representative images of WT (CBY8089) and Δ-s-tether cells (CBY8091) and nuclei showing expression of genomically-integrated Cdc5-GFP at different stages through the cell cycle; Cdc5 expression levels were highly variable in Δ-s-tether cells. Arrowheads indicate Cdc5 at the bud neck, which is observed earlier in Δ-s-tether cells in medium budded cells. Scale bar: 10 µm. **(D)** Quantification of Cdc5-GFP in WT and Δ-s-tether cells at different cell-cycle stages. Percentage of medium and large budded cells exhibiting normal and defective Cdc5 localizations as depicted. \*\*\**p* = 0.0001; n = 50 cells for each morphological class. **(E)** Model illustrating pathways that integrate cER-PM MCSs and phospholipid regulation with cell-cycle control during late mitosis. In this context, Elm1 phosphorylates and activates Hsl1, independent of their regulation of septin assembly. In turn, activated Hsl1 phosphorylates and inhibits Pah1. To maintain DAG:PA balance, Pah1 counteracts Dgk1 synthesis of PA and cER-PM MCSs regulation of phospholipid flux. We propose that Elm1 “senses” cER-PM MCSs through its monitoring of cellular DAG:PA or cortical actin polarization (as a proxy for nucleopodia), or both. As an alternative possibility, because of nuclear efflux through membrane ruptures, Cdc5 spatiotemporal misregulation might interfere with Elm1 when they are both recruited to the bud neck. Proteins shown in green letters indicate positive regulators of cER-PM MCSs (as determined through Δ-s-tether genetic interactions), whereas those shown in red indicate proven negative regulators and effectors. Proteins in bold lettering did not exhibit any functional interactions when eliminated from Δ-s-tether cells, and the underlined Cdc5 indicates its spatiotemporal regulation is perturbed by the loss of cER-PM MCS.

In addition to its role in SPOC signaling, the nonessential Hsl1p kinase is a Pah1 inhibitory kinase, which we hypothesized might functionally connect the regulation of Pah1p activity with the observed cell-cycle delay in Δ-s-tether cells. Compared to WT cells, *hsl1*Δ cells have no growth defect at any temperature. Given that *PAH1* deletion suppresses Δ-s-tether growth defects, we predicted that deletion of a negative regulator of Pah1 would have the opposite effect; accordingly, *HSL1* deletion in Δ-s-tether cells is lethal at 37°C and severely impairs growth at 30°C (Fig 4A). Choline partially rescues the *hsl1*Δ Δ-s-tether growth defects, suggesting that phospholipid regulation underlies the functional interaction between *HSL1* and cER-PM MCSs (Fig 4A). As an upstream Hsl1 activator that also plays an important role in SPOC regulation, the Elm1 kinase is also essential for viability in Δ-s-tether cells, in which the deletion of *ELM1* in Δ-s-tether cells is lethal even with choline supplementation (Fig 6B). Because choline otherwise suppresses Δ-s-tether defects, we tested if *elm1*Δ Δ-s-tether lethality is contingent on the physical reestablishment of cER-PM contact as opposed to phospholipid flux. Choline supplementation was insufficient to suppress *elm1*Δ Δ-s-tether lethality, though *DGK1* overexpression restored viability. In contrast, the artificial staple alone did not rescue *elm1*Δ Δ-s-tether inviability, but the combination of choline and the artificial staple supported slow growth (S4 Fig). These results suggest that in cells lacking cER-PM tethers, Elm1 is required to maintain the DAG:PA balance at least partly through Hsl1. This balance can be partially restored by nonspecific reestablishment of cER-PM contact, but only when combined with increased choline-dependent phospholipid production; suggesting the physical contact conferred by cER-PM MCSs partially contributes to Elm1 regulation (S4 Fig). Although Elm1 is necessary for the assembly of septin subunits at the bud neck, this activity seems irrelevant in Δ-s-tether cells given that Cdc12-GFP septin organization appears unaffected [90–94] (Fig 5E). Moreover, other septin-associated genes including the anillin *BUD4* and the nonessential septin *SHS1*, both implicated in nuclear segregation defects in other yeasts [87], do not impact growth when deleted in Δ-s-tether cells (S1 Table). In cells lacking cER-PM contact, SPOC appears to monitor nuclear membrane lipids, rather than responding to microtubules or septin changes in the bud neck.

The Protein Phosphatase 2A (PP2A) holoenzyme contains either Cdc55 or Rts1 as its regulatory subunit, and each isoform performs distinct roles in controlling PP2A activity during mitosis [95]. In SPOC signaling, PP2A^Rts1^ dephosphorylates the Kin4 kinase, which directs Kin4 localization to the yeast microtubule organizing center/SPB and its function in SPOC activation [96]. Neither *RTS1* nor *KIN4* deletion significantly alters Δ-s-tether growth, but deletion of *CDC55* is lethal in these cells (S1 Table) (S4 Fig). PP2A^Cdc55^ activity intersects with SPOC signaling pathway at multiple points and contributes to mitotic entry through regulation of the morphogenesis checkpoint effector Swe1 [51, 97]. However, deletion of *SWE1* in Δ-s-tether cells does not exacerbate the growth defect, suggesting that PP2A^Cdc55^ activity essential in Δ-s-tether cells is independent of Swe1 (Fig 6A). Notably, Elm1, Hsl1, and Cdc55 localize to the bud neck during late mitosis, raising the possibility that this region of the PM plays a significant role in the cell-cycle delay observed in cells lacking cER-PM MCSs. Alternatively, the functional interaction between cER-PM tethers and PP2A^Cdc55^ might reflect another non-canonical PP2A^Cdc55^ activity [98].

As a key effector of SPOC signaling, the yeast polo-like kinase Cdc5 plays an important role in nuclear morphological and nuclear envelope dynamics changes during mitosis [47, 48, 92, 99–101]. Because Cdc5 distribution during mitosis reflects its distinct activities inside and outside the nucleus, we compared Cdc5-GFP localization in WT and Δ-s-tether cells to test whether cER-PM MCSs alter Cdc5 spatiotemporal regulation. In WT cells, Cdc5 translocates from the SPB into the nucleus where it accumulates during S-phase. Later during anaphase, Cdc5 is released into the cytoplasm where it interacts with the bud neck and SPBs (Fig 6C and 6D) [52]. In Δ-s-tether cells, however, there is a significant increase in Cdc5-GFP fluorescence at the bud neck in medium-budded cells (Fig 6C and 6D). Moreover, while Cdc5-GFP typically localizes within the nucleus of budded cells, distinct Cdc5-GFP fluorescence can be observed at the nuclear membrane in 7.8% of Δ-s-tether cells (n = 235 cells; *p* = 0.002). Supernumerary Cdc5-GFP fluorescent SPBs are also detected in a small but significant subset of Δ-s-tether cells, consistent with the multiploidy driven by DNA duplication without concomitant karyokinesis, yielding some nuclei with 3 visible SPBs (4.7%; n = 244 cells; *p* = 0.006). Although the *CDC5*-GFP construct is integrated at its native site and expressed under its native promoter, it should be noted that in Δ-s-tether cells *CDC5*-GFP fluorescence intensity is variable. The variability in Cdc5 expression does not appear to be linked to transcriptional changes, given that *CDC5* transcription remains unchanged in Δ-s-tether cells relative to WT, as previously determined through transcriptomics [15]. In terms of post-transcriptional regulation, Cdc5 levels, like other mitotic regulators, are reduced through Cdc14-and Cdh1-dependent degradation by anaphase-promoting complex/cyclosome complex (APC/C^Cdh1^) [102–107]. In Δ-s-tether cells, however, neither *CDH1* deletion nor overexpression affects growth (S1 Table), suggesting that APC/C^Cdh1^–dependent changes in Cdc5 levels are not a contributing factor. Nevertheless, Cdc5 spatiotemporal dysregulation in these cells provides an additional read-out of nuclear disruptions associated with the SPOC-induced delay.

## Discussion

Our analysis supports a model in which yeast cER-PM MCSs reinforce nuclear membrane integrity through two independent mechanisms. First, they physically anchor the nucleus to the cell cortex, which is predicted to dampen nuclear membrane torsion. In this way, we propose that linking the PM to the nucleus through cER extensions acts like mooring lines, structurally supporting the nuclear envelope to prevent membrane tearing and enable proper positioning of the nucleus relative to the plane of cell division. Second, cER-PM MCSs coordinate of ER-PM phospholipid biosynthesis to modulate nuclear ER composition for maintaining bilayer cohesion [13, 108]. In cells lacking cER-PM MCSs, the combined loss of external attachment and altered membrane lipid composition results in severe nuclear ER rupturing that exceeds defects observed in established mutants compromised in nuclear membrane repair (e.g. *chm7*Δ). Although Scs2 represents the major tether contributing mostly to cER-PM contact [9], and *SCS2* deletion directly affects cER dynamics [55], nuclear envelope compartmentalization was compromised only when all cER-PM tether deletions were combined, consistent with the observed growth inhibition. The redundancy of these external and internal nuclear mechanisms for maintaining the nuclear ER is further demonstrated by the two independent modes of Δ-s-tether suppression: restoration of non-specific cER-PM contact via artificial staple, and alteration of nuclear ER membrane lipid distribution (i.e. choline addition, *PAH1* deletion, or *DGK1* overexpression). These mechanisms act independently given that the artificial staple confers limited cER-PM contact but does not restore normal phospholipid synthesis, whereas *DGK1* overexpression rescues phospholipid defects in cells lacking cER-PM tethers but does not restore cER-PM association [14, 16]. The nuclear integrity defects that result from the removal of ER-PM MCSs are tolerated due to a robust ESCRT-III nuclear membrane repair mechanism. Loss of proteins involved in either nuclear ESCRT-III-mediated membrane repair, or by removing the extra repair time provided via the SPOC checkpoint cell-cycle delay, significantly exacerbates growth defects of cells lacking cER-PM MCSs. Unlike extreme nuclear lesions caused by specific laminopathies and chromothripsis [109], we show that simple UPR induction by ER stress does not disrupt nuclear envelope containment. These results are consistent with previous findings indicating that UPR induction only affects the expansion of the perinuclear space without affecting membrane integrity [110]. Beyond yeast, these findings have implications for the maintenance of nuclear integrity and genomic stability in metazoan cells, as well as for maintaining membrane homeostasis of other organelles. It has been shown in mammalian cells, that tension arising from cER-PM tethering propagates mechanotransduction to other membranes, which impacts general cellular metabolism [111].

In yeast, the PM is associated, on average, with at least fivefold more cER than in mammalian cells, which might reflect differences in how these cells maintain nuclear ER membrane structure [13]. In metazoans, the nuclear lamins provide a structural scaffold that further buttresses the inner nuclear membrane, and they interact with LEM (LAP2-Emerin-MAN1) domain proteins to support nuclear stability. Although yeast lack nuclear lamins, they do contain a LEM homologue Heh1, which is also a key adaptor for ESCRT-III nuclear membrane repair, and is essential in Δ-s-tether cells [66, 112–115]. Intralumenal nuclear membrane linkages between SUN/KASH domain proteins also lend additional support to inner and outer nuclear membranes [116–118]. However, while yeast SUN/KASH-like protein complexes are important in SPB assembly, their role in maintaining overall nuclear structure is less clear [119, 120]. We propose that, in lieu of the internal structural support provided by the nuclear lamina and SUN/KASH trans-nuclear membrane connectors in metazoan cells, the nuclear envelope in yeast is partially stabilized by external support provided by cER-PM contact. Although less extensive than in yeast, cER-PM association in some mammalian cell types is substantial, which might contribute to nuclear envelope stability. For example, in neuron cell bodies, 12.5% of the inner PM surface is covered by cER, suggesting that in those cells cER-PM MCSs might play an important role in maintaining nuclear integrity [121, 122]. Under specific conditions, such as during acute mechanical stresses on the nucleus including that associated with cell migration through confined spaces, cER-PM tethering might provide necessary support for stabilizing the nuclear membrane.

Inhibition of sphingolipid biosynthesis in budding yeast has been reported to reduce nuclear volume and causes irregular nuclear morphology [123]. In addition to reduced phospholipid flux in Δ-s-tether cells, lipidomic analyses also revealed a significant decrease in sphingolipids levels upon deletion of cER-PM tethers [14, 16]. In Δ-s-tether cells, however, sphingolipid dysregulation appears to extend beyond a reduction in synthesis as these cells are resistant (rather than sensitive, as might otherwise be predicted) to lethal concentrations of the sphingolipid synthesis inhibitor myriocin [14]. Regardless, unlike sphingolipid mutants, overall size and morphology of nuclei in Δ-s-tether cells do not differ significantly from WT. Furthermore, unlike cells lacking cER-PM MCSs, sphingolipid depletion has not been linked to nuclear membrane rupturing, induction of ER stress pathways, or cell-cycle delays [9, 14, 15, 123]. Even in immortalized human cell lines, sphingolipid depletion fails to trigger a cell cycle delay despite enhancing chromothripsis-like genetic instability [123]. Furthermore, sphingolipids and phospholipids appear to affect nuclear membrane expansion in different ways. Somewhat similar to sphingolipid-depleted cells, *pah1*Δ cells exhibit irregular nuclear morphology, but their nuclei are not reported to be smaller [21]. Unlike sphingolipid-or *PAH1*-associated nuclear defects, cells lacking cER-PM MCSs exhibit nuclear membrane blisters and ruptures but maintain a nuclear roundness comparable to WT. Taken together, these differences indicate that the nuclear defects in cells lacking cER-PM MCSs differ from those caused by sphingolipid synthesis inhibition, which are themselves distinct from those in *PAH1* deletion cells.

The proper balance of DAG and PA levels within the nuclear envelope play an important role in maintaining membrane morphology and integrity [74]. When *DGK1* is overexpressed or in *pah1*Δ mutants, DAG levels are reduced and PA levels increase, the opposite of the lipid imbalance observed in Δ-s-tether cells by lipidomics [14, 16, 21]. The restoration of the lipid balance likely explains why *DGK1* overexpression and *pah1*Δ suppress Δ-s-tether defects. We next considered how these lipid changes specifically impact the nuclear membrane. In Δ-s-tether cells, DAG levels rise in nuclear membranes, consistent with the overall increase in cellular DAG levels, and highly motile intranuclear DAG-enriched lipid aggregates accumulate. Counterintuitively, nuclear PA levels are not reduced but instead increase. Even though both DAG and PA are enriched in Δ-s-tether nuclei, the nuclear PA:DAG ratio is still reduced compared to WT, indicating a substantial perturbation in both the PA-DAG balance and the absolute levels of these lipids. These findings suggest that the relative balance and, potentially, the absolute levels of these two lipids contribute to the maintaining nuclear membrane integrity in cells lacking ER-PM MCSs.

Despite nuclear segregation defects that lead to an accumulation of multiploid cells and extensive nuclear membrane rupture, deletion of cER-PM tethers does not cause chromosome instability or trigger a SAC response. This result contrasts with the extensive chromosomal disruptions evident during chromothripsis in cancer cells following nuclear membrane disruption, which can expose nuclear DNA to N4BP2 and TREX1 cytoplasmic nucleases, neither of which is present in yeast [124–126]. However, the efflux of DNA repair enzymes during nuclear membrane disruption has been proposed to increase spontaneous mutation rates in cancer cells [4, 76]. Consistent with this idea, nuclear ruptures in Δ-s-tether cells appear to increase the spontaneous mutation frequency, supporting the model that nuclear envelope maintenance affects DNA damage surveillance in yeast [83].

Apart from nuclear envelope rupturing, our analysis also suggests that nuclear support and positioning by cER-PM contacts contribute to the fidelity of nuclear segregation and division during yeast closed mitosis. In budding yeast, previously reported mutants that disrupt nuclear segregation typically exhibit defects either in astral microtubules/SPB attachments or in septin organization at the bud neck [46, 127, 128]. Microtubules or septins, however, are not significantly affected by the elimination of ER-PM MCSs. To our surprise, cortical actin patches show marked depolarization throughout the cell cycle in Δ-s-tether cells, though this defect is insufficient to interfere with bud formation. Nucleopodia represent microtubule-independent nuclear envelope protrusions that initiate membrane remodeling to assist the complete migration of the anaphase nucleus through the bud [44]. Consistent with the actin-and phospholipid-dependence of nucleopodial formation, nucleopodia formation was inhibited in cells lacking cER-PM MCSs. This defect is distinct from other classes of mutants in which karyokinesis is unaffected but segregation of the daughter nucleus into the bud fails, leading to the accumulation of binucleated cells [46]. In cells lacking cER-PM MCSs, a unique conjunction of defects in preanaphase nuclear positioning, actin depolarization, and changes in DAG:PA levels appear to elicit a SPOC-dependent cell-cycle delay that persists until membrane remodeling and dynamics are restored.

Consistent with a SPOC-dependent cell-cycle delay when cER-PM tethers are absent, deletion of several SPOC regulatory genes in Δ-s-tether cells results in lethality or significant growth defects, indicating strong genetic interactions. Most notably, the synthetic lethality of *ELM1* deletion and the synthetic growth defect of *HSL1* deletion in Δ-s-tether cells point to a functional interaction connecting phospholipid regulation and SPOC signaling. In addition to its role in SPOC signaling, the Hsl1 kinase inhibits Pah1 generation of DAG, a function that parallels the regulatory activity of cER-PM MCSs on phospholipid flux (Fig 6E). Because *ELM1* is an upstream activator of *HSL1*, yet deletion of *ELM1* deletion in Δ-s-tether cells causes a more severe growth defect than deletion of *HSL1*, Elm1 appears to respond to additional aspects of cER-PM contact beyond phospholipid levels or distribution. The polo-like kinase Cdc5 integrates nuclear events during the cell cycle and, though it has not yet been reported to regulate Elm1, the functions of both kinases intersect at the bud neck. In Δ-s-tether cells, variability in the visualization of Cdc5-GFP might be a consequence of nuclear envelope rupture that in turn permits Cdc5 premature association with the bud neck. Such spatiotemporal misregulation of Cdc5 is consistent with a model in which Cdc5 interferes at the bud neck with Elm1-dependent regulation of SPOC and Hsl1-mediated control of phospholipid flux (Fig 6E). Alternatively, Elm1 acts as a coincidence detector at the bud neck, sensing defects in cortical actin patches (hence nucleopodia formation) and the cellular balance of DAG:PA, while Cdc5 variability and spatiotemporal dysregulation might simply be a downstream consequence of nuclear membrane instability.

## Materials and Methods

### Strains and Plasmids

Yeast strains used are listed in the S1 Table and plasmids in S2 Table. DNA cloning and bacterial and yeast transformations were carried out using standard techniques as previously described [14]. Unless otherwise stated, yeast cultures were grown in synthetic minimal or YPD rich media at 30°C. For the choline supplementation assays, yeast strains were cultured in synthetic minimal media for 48 h prior to analysis or streaked onto solid synthetic complete media containing 1 mM choline chloride (Sigma-Aldrich Chemicals, St. Louis, MO). The inherent genetic instability of Δ-s-tether cells requires that the strain is maintained by culturing at 30°C on synthetic medium supplemented with 1 mM choline and/or transformed with a *URA3*-marked plasmid containing *SCS2*. Before analysis of Δ-s-tether cells in assays or for other molecular manipulations, the *SCS2* plasmid is selected against by growth on synthetic complete solid growth medium containing 1 g/L 5’-fluoroorotic acid (FOA), after which they are then evaluated.

The methodology for constructing these CRISPR-Cas9 plasmids was as previously described [129], and successful deletions were confirmed by PCR. Because of the genetic instability in cells lacking cER-PM tethers, which interferes with genomic integration, additional gene deletions were generated in Δ-s-tether cells expressing episomal *SCS2*, which complements the membrane tethering defect. Furthermore, to promote efficient gene deletion, these cells were transformed with specific CRISPR-Cas9 plasmids. To evaluate the effect of CRISPR-Cas9 induced mutations in Δ-s-tether cells, for the analysis of genetic interactions, cells were treated with 5’-FOA to select for the absence of the *URA3*-marked *SCS2*-containing plasmid.

The yeast GFP-cGAS expressing plasmid pCB1465 was constructed by amplifying the GFP-cGAS fragment by polymerase chain reaction (PCR) from pCB1424. The amplified product was inserted into the EcoRI/BamHI restriction sites of pCB380, which generated a GFP-cGAS gene fusion with the *ACT1* promoter for its constitutive expression in yeast. The yeast Chm7–mRuby2 expression plasmid (pCB1661) was constructed using the Golden Gate cloning system in combination with the MoClo yeast toolkit in which the *CHM7* coding sequence, including its endogenous promoter but lacking the stop codon and 3ʹ UTR, was PCR-amplified and fused with mRuby2 coding sequence [130].

### Electron microscopy

For cryofixation and electron microscopy, yeast cells were grown to mid-log phase at 30°C and concentrated in 200 µl of growth media. Cell suspensions were cryo-preserved using EM ICE High Pressure Freezer (Leica Mikrosysteme, Vienna, Austria). Vitrified samples underwent water substitution in a Leica EM ASF2 freeze substitution system for 3 days in 0.1% uranyl acetate and 1% osmium tetroxide in acetone. Temperature was gradually increased from -90°C to -70°C (20 h), from -70°C to -50°C (16 h), from -50°C to -20°C (20 h) and finally from -20°C to 20°C (5 h). Stained samples were washed with acetone and infiltrated with Spurr’s low viscosity embedding resin (Electron Microscopy Sciences, Hartfield, USA) and hardened at 60°C. Ultra-thin sections were stained with 2% uranyl acetate for 6 min and Reynold’s lead citrate for 3 min. Sections were imaged on a FEI Tecnai Spirit TEM at 120kV (Thermo Fisher Scientific, Ottawa, Canada). The nuclear envelope width for a single nucleus represents the mean of 5 measurements collected from randomly selected locations along the nuclear membrane; each measurement of nuclear envelope width includes both bilayers of the inner and outer nuclear membranes and the perinuclear space. For quantification of intranuclear lipid droplet aggregates, nucleoplasmic structures devoid of negative stain were counted if their diameter exceeded ∼30 nm. Digital images were processed and colourized using Affinity Photo software (Nottingham, UK). TEM images represent triplicate biological replicates.

### Flow cytometry

Mid-log yeast cultures were prepared as previously described [131] and analyzed on a BD LSRFortessa X-20 Cell Analyzer (BD Biociences, Mississauga, Canada) to measure DNA content. Prior to quantification, background signal and doublets were removed. Cells were quantified by gating 1C, 2C and 4C DNA content. Quantification of DNA content in each strain represents three independent trials.

### Fluorescence microscopy

Wide-field fluorescence imaging was performed as previously described, with modifications, using a Leica DMXRA2 microscope equipped with a 100/1.40 Plan-Apo objective; images were acquired with an Orca100ER or AG digital camera (Hamamatsu Photonics, Hamamatsu-City, Japan) [132]. Unless otherwise noted, exposure times and contrast enhancement were constant for a given series of images. To examine nuclear DNA staining, mid-log phase cells were pelleted and resuspended in 1 mL 3:1 (v/v) methanol/acetic acid. After 30 min at room temperature, cell pellets were resuspended in 1 mL PBS. Following an additional wash with 1 mL PBS, the final pellet was resuspended in mounting medium containing 1 mg/mL 4’,6-diamidino-2-phenylindole (DAPI; Accurate Chemicals and Scientific Corp., Westbury, NY, USA) for nuclear fluorescence staining. DAPI nuclear fluorescence was captured with 50 ms exposures. For nuclear rupture quantification, mid-log phased yeast cells expressing GFP-cGAS from the *ACT1* promoter a low-copy plasmid (pCB1465) were cultured at 30°C and imaged with 100 ms exposures at a gain of 225. GFP-cGAS localization was defined as nuclear when the mean nuclear-to-cytoplasmic fluorescent intensity ratio exceeded 1.5. To examine septins and tubulin, Cdc12-GFP and Venus- Tub1, respectively, were visualized in live log-phased cells cultured at 30°C and imaged with a 100 ms exposure at a gain of 255. To examine nuclear PA and DAG, cells expressing NLS-DAG-GFP and NLS-PA-mCherry, respectively, were cultured to mid-log phase at 30°C. NLS-PA-mCherry images were acquired using a 100 ms exposure time. NLS-DAG-GFP was imaged using exposure times of 50, 100 and 200 ms depending on strain. For quantitative analysis, NLS-DAG-GFP fluorescence intensity was compared from images acquired at 100 ms exposures following Fast Restoration deconvolution. Nuclear fluorescence quantification of NLS-PA-mCherry included the entire nucleus whereas quantification for NLS-DAG-GFP was restricted to just the nuclear envelope, without nucleoplasm (thereby excluding most intranuclear aggregates). To examine the spatiotemporal localization of Cdc5, GFP was integrated at the 3’-end of *CDC5* and, after culturing at 30°C to mid-log phase, live cells were imaged with 500 ms exposures at a gain of 255. To quantify Chm7-mRuby2 recruitment to nuclear ruptures, yeast cells coexpressing GFP-cGAS and Chm7-mRuby2 were cultured at 30°C to mid-log phase and imaged after a 100 ms exposure for GFP-cGAS and 500 ms for Chm7-mRuby2. Analysis of all widefield microscopy images was conducted using Volocity 7.0.0 imaging software (Quorum Technologies Inc., Puslinch, ON).

Images captured by super resolution microscopy were acquired on a Zeiss LSM 880 confocal microscope with an Airyscan fluorescence GaAsP detector and 63x/1.4 oil immersion objective (Carl Zeiss Canada Ltd., Toronto, ON). To analyze cortical actin polarization, cells were fixed and stained with rhodamine conjugated phalloidin as previously reported [133]. Rhodamine-phalloidin fluorescence was imaged using the 561 nm laser set at 1.5% laser intensity with the detector gain set at 900; the pixel dwell time was 1.37 μs for 11 z-stacks through 1.8 μm sections. Images were processed using Airyscan joint deconvolution (5 iterations at a 95% confidence limit) in ZEN black software, which was then used to quantify relative fluorescence polarization. Cortical actin patch polarization is defined as the ratio of cortical fluorescence pixel density in polarized regions relative to that along the rest of the cell cortex. For super-resolution imaging of Chm7-mRuby2 recruitment to GFP-cGAS nuclear ruptures, mRuby2 and GFP were excited using the 561 nm (at 1.5% power) and the 488 nm laser set at (2% power) lasers, respectively. For both channels, pixel dwell time was 5.25 μs for 15 z-stacks acquired through 5.6 μm sections, with the detector gain at 800.

### Time lapse fluorescence microscopy

Time lapse fluorescence microscopy for GFP-cGAS nuclear rupturing was performed by super-resolution fluorescence microscopy on a Zeiss LSM 880 confocal microscope with an Airyscan super-resolution GaAsP detector and 63x/1.4 oil immersion objective. Cells were cultured to exponential phase after which they were continuously imaged at 30°C in synthetic complete media for 1.5 hrs. GFP-cGAS fluorophore imaged with the 488 nm laser (2% power) and digital gain set at 800; dwell times were 1.19 μs per pixel imaging 11 Z-stacks through 9 μm sections. The time lapse Airyscan images were processed in ZEN Black prior to analysis and each GFP-cGAS time series represents 6 images taken at 15 min intervals.

The 4D analysis of nucleopodia was conducted in asynchronous log-phased cells co-expressing Venus-Tub1 and the NLS-PA-mCherry nuclear probe (pRS316-NLS-Opi1-Q2-mCherry) cultured in synthetic complete liquid media at 30°C. Time lapse fluorescence microscopy was performed on a Zeiss LSM 880 confocal microscope with an Airyscan super-resolution GaAsP detector and 63x/1.4 oil immersion objective with a dwell time of 1.54 μs per pixel. NLS-PA-mCherry fluorophore was excited using 561nm laser set at 2.5% laser intensity and digital gain set at 900. Venus-Tub1 was excited using 514nm laser set at 3% laser intensity and the digital gain set at 750. The time-lapse images were Airyscan processed in ZEN Black, and the time series represents images taken at 30 s intervals.

Videos of highly mobile intranuclear DAG particles were obtained by wide-field microscopy using a Leica DMXRA2 microscope equipped with a 100x/1.40 Plan-Apo objective and images were processed using Volocity 7.0.0 software. During imaging, mid-log cells coexpressing NLS-DAG-GFP and NLS-PA-mCherry were cultured at 30°C in synthetic complete liquid medium. Image time series were as described. All time-lapse videos represent ≥ 3 independent trials.

### Genomic instability and fluctuation assays

Plasmid-loss chromosomal instability assays were adapted with modifications from previous protocols [134, 135]. Yeast strains transformed with a *URA3*-marked *CEN* plasmid were cultured overnight in selective medium lacking uracil, diluted in non-selective rich liquid medium to equivalent cell densities and were cultured at 30 °C. To yield ∼600 colonies on solid non-selective growth plates, log-phased cells were again diluted before plating and then grown for 3 days at 30 °C. To identify colonies in which the plasmid was retained without selection, and to quantify those that did not, nonselective growth plates were replica-printed onto solid medium selecting for the *CEN* plasmid and incubated at 30°C for 2 days; plasmid loss is indicated by the relative percentage of colonies that can only grow on non-selective (indicating that the *URA3 CEN*-plasmid was not retained). Three biological replicates were performed for each yeast strain.

Fluctuation assays were performed with modifications as previously reported [84, 136]. Single colonies were inoculated into synthetic liquid medium lacking uracil and leucine, to maintain *URA3* and *LEU2* prototrophies of the *CEN* plasmid (pCB1774) transformants. Cultures were grown overnight at 30°C, then diluted into fresh synthetic complete medium to a density of 4000 cells/ml. Cells were subsequently aliquoted into multiwell plates at 100 µL per well and cultured at 30°C for 2-3 days to culture saturation. To determine the total number of plasmid transformants (with and without spontaneous *ura3* mutations), 8 wells were randomly selected, serially diluted for counting, and plated on solid synthetic medium lacking only leucine. To count 5-FOA-resistant *ura3* mutant transformants, the remaining 89 cultures were spotted onto synthetic medium lacking leucine and containing 0.1% 5-FOA. After incubation at 30°C for 3 days, spontaneous 5-FOA-resistant *ura3* mutant colonies were counted for each independent spotted culture.

### Statistical Analysis

Spontaneous mutation frequencies from fluctuation tests were estimated from colony counts on selective and nonselective plates using the rSalvador R package [137]. Pairwise likelihood ratios were used to calculate 95% confidence intervals and determine *p*-value statistical differences in relative mutation frequencies [137].

Statistical analyses on binary categorical data used Fisher’s exact test, otherwise unpaired student t tests were used. In all graphs, error bars represent standard deviations. All graphs represent n ≥ 3 independent trials.

## Acknowledgements

Many thanks to Tim Audas, Dheva Setiaputra, and Peter Stirling for helpful discussions and comments about this work. The authors thank the generosity of Scott Emr, Philip Hieter, Alwin Köhler, Christopher Loewen, Peter Novick and Chris Stefan for sharing plasmids and yeast strains.

## Supporting Information

**S1 Table.**
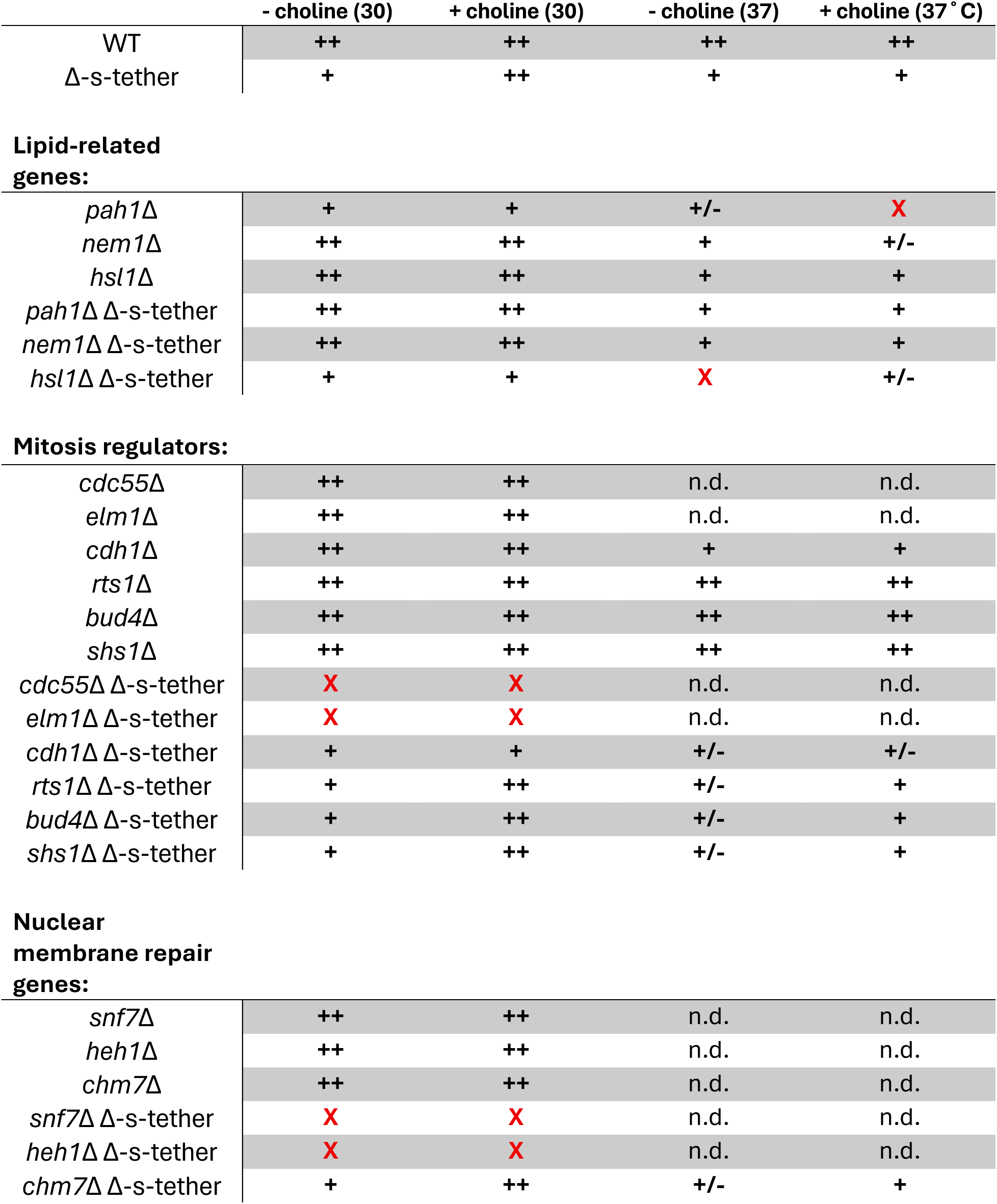
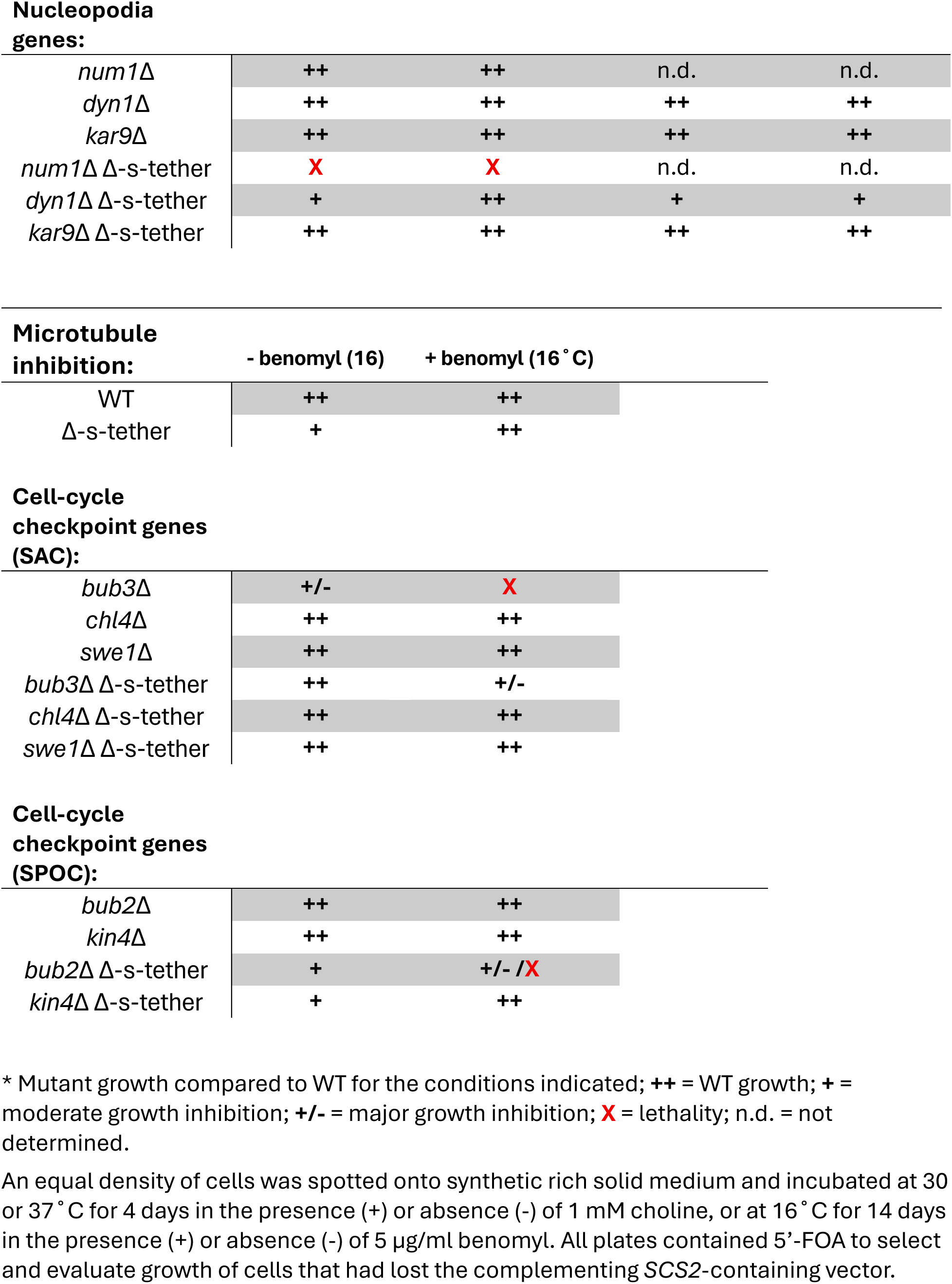
. Summary of genetic interactions.

|  | **- choline (30)** | **+ choline (30)** | **- choline (37)** | **+ choline (37˚C)** |
|---|---|---|---|---|
| WT | **++** | **++** | **++** | **++** |
| Δ-s-tether | **+** | **++** | **+** | **+** |
| **Lipid-related genes:** |  |  |  |  |
| *pah1Δ* | **+** | **+** | **+/-** | **X** |
| *nem1Δ* | **++** | **++** | **+** | **+/-** |
| *hsl1Δ* | **++** | **++** | **+** | **+** |
| *pah1Δ* Δ-s-tether | **++** | **++** | **+** | **+** |
| *nem1Δ* Δ-s-tether | **++** | **++** | **+** | **+** |
| *hsl1Δ* Δ-s-tether | **+** | **+** | **X** | **+/-** |
| **Mitosis regulators:** |  |  |  |  |
| *cdc55Δ* | **++** | **++** | n.d. | n.d. |
| *elm1Δ* | **++** | **++** | n.d. | n.d. |
| *cdh1Δ* | **++** | **++** | **+** | **+** |
| *rts1Δ* | **++** | **++** | **++** | **++** |
| *bud4Δ* | **++** | **++** | **++** | **++** |
| *shs1Δ* | **++** | **++** | **++** | **++** |
| *cdc55Δ* Δ-s-tether | **X** | **X** | n.d. | n.d. |
| *elm1Δ* Δ-s-tether | **X** | **X** | n.d. | n.d. |
| *cdh1Δ* Δ-s-tether | **+** | **+** | **+/-** | **+/-** |
| *rts1Δ* Δ-s-tether | **+** | **++** | **+/-** | **+** |
| *bud4Δ* Δ-s-tether | **+** | **++** | **+/-** | **+** |
| *shs1Δ* Δ-s-tether | **+** | **++** | **+/-** | **+** |
| **Nuclear membrane repair genes:** |  |  |  |  |
| *snf7Δ* | **++** | **++** | n.d. | n.d. |
| *heh1Δ* | **++** | **++** | n.d. | n.d. |
| *chm7Δ* | **++** | **++** | n.d. | n.d. |
| *snf7Δ* Δ-s-tether | **X** | **X** | n.d. | n.d. |
| *heh1Δ* Δ-s-tether | **X** | **X** | n.d. | n.d. |
| *chm7Δ* Δ-s-tether | **+** | **++** | **+/-** | **+** |
| **Nucleopodia genes:** |  |  |  |  |
| *num1Δ* | **++** | **++** | n.d. | n.d. |
| *dyn1Δ* | **++** | **++** | **++** | **++** |
| *kar9Δ* | **++** | **++** | **++** | **++** |
| *num1Δ* Δ-s-tether | **X** | **X** | n.d. | n.d. |
| *dyn1Δ* Δ-s-tether | **+** | **++** | **+** | **+** |
| *kar9Δ* Δ-s-tether | **++** | **++** | **++** | **++** |
| **Microtubule inhibition:** | **- benomyl (16)** | **+ benomyl (16˚C)** |  |  |
| WT | **++** | **++** |  |  |
| Δ-s-tether | **+** | **++** |  |  |
| **Cell-cycle checkpoint genes (SAC):** |  |  |  |  |
| *bub3Δ* | **+/-** | **X** |  |  |
| *chl4Δ* | **++** | **++** |  |  |
| *swe1Δ* | **++** | **++** |  |  |
| *bub3Δ* Δ-s-tether | **++** | **+/-** |  |  |
| *chl4Δ* Δ-s-tether | **++** | **++** |  |  |
| *swe1Δ* Δ-s-tether | **++** | **++** |  |  |
| **Cell-cycle checkpoint genes (SPOC):** |  |  |  |  |
| *bub2Δ* | **++** | **++** |  |  |
| *kin4Δ* | **++** | **++** |  |  |
| *bub2Δ* Δ-s-tether | **+** | **+/- /X** |  |  |
| *kin4Δ* Δ-s-tether | **+** | **++** |  |  |

\* Mutant growth compared to WT for the conditions indicated; **++** = WT growth; **+** = moderate growth inhibition; **+/-** = major growth inhibition; **X** = lethality; n.d. = not determined.
An equal density of cells was spotted onto synthetic rich solid medium and incubated at 30 or 37˚C for 4 days in the presence (+) or absence (-) of 1 mM choline, or at 16˚C for 14 days in the presence (+) or absence (-) of 5 µg/ml benomyl. All plates contained 5’-FOA to select and evaluate growth of cells that had lost the complementing *SCS2*-containing vector.

**S2 Table.**
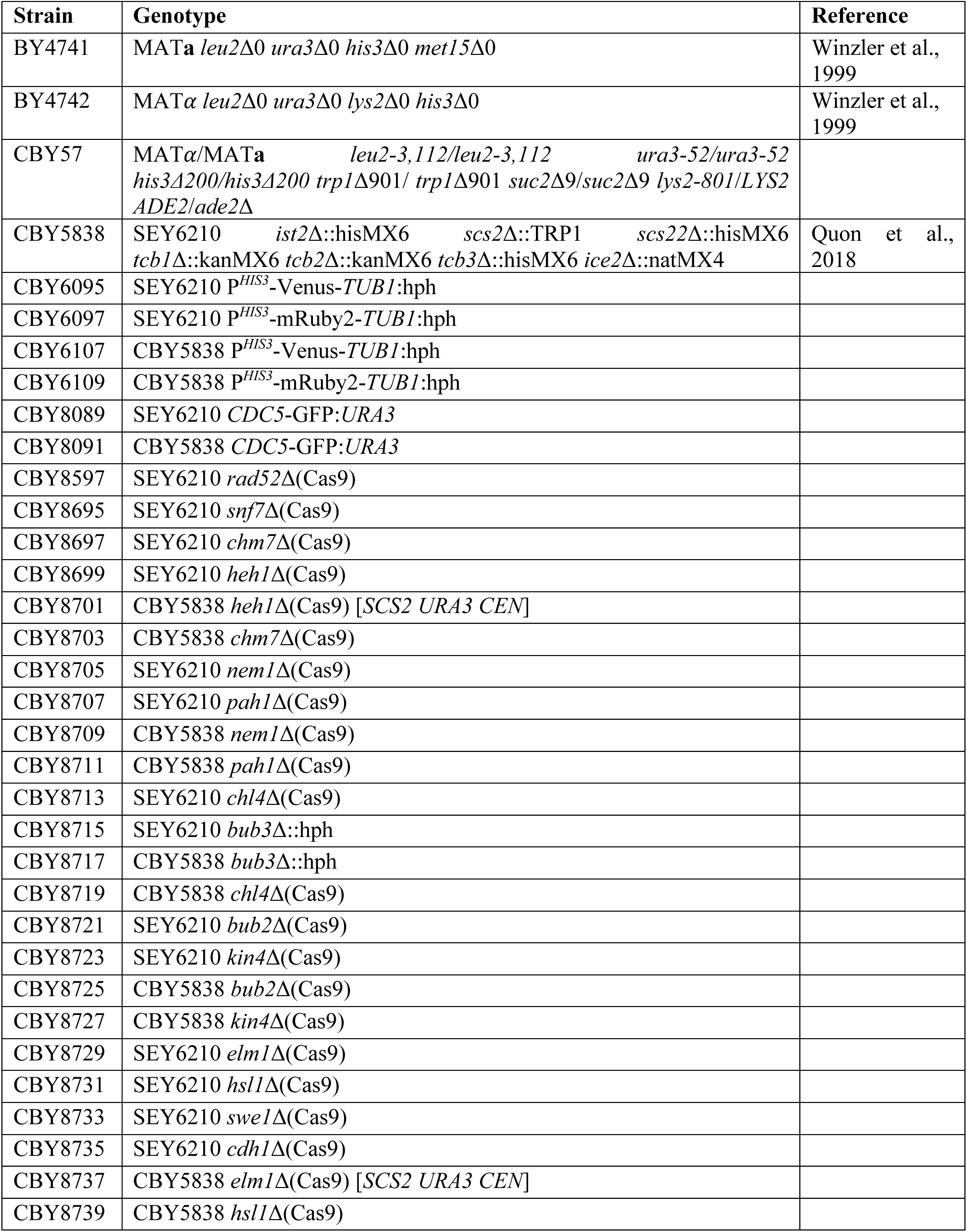
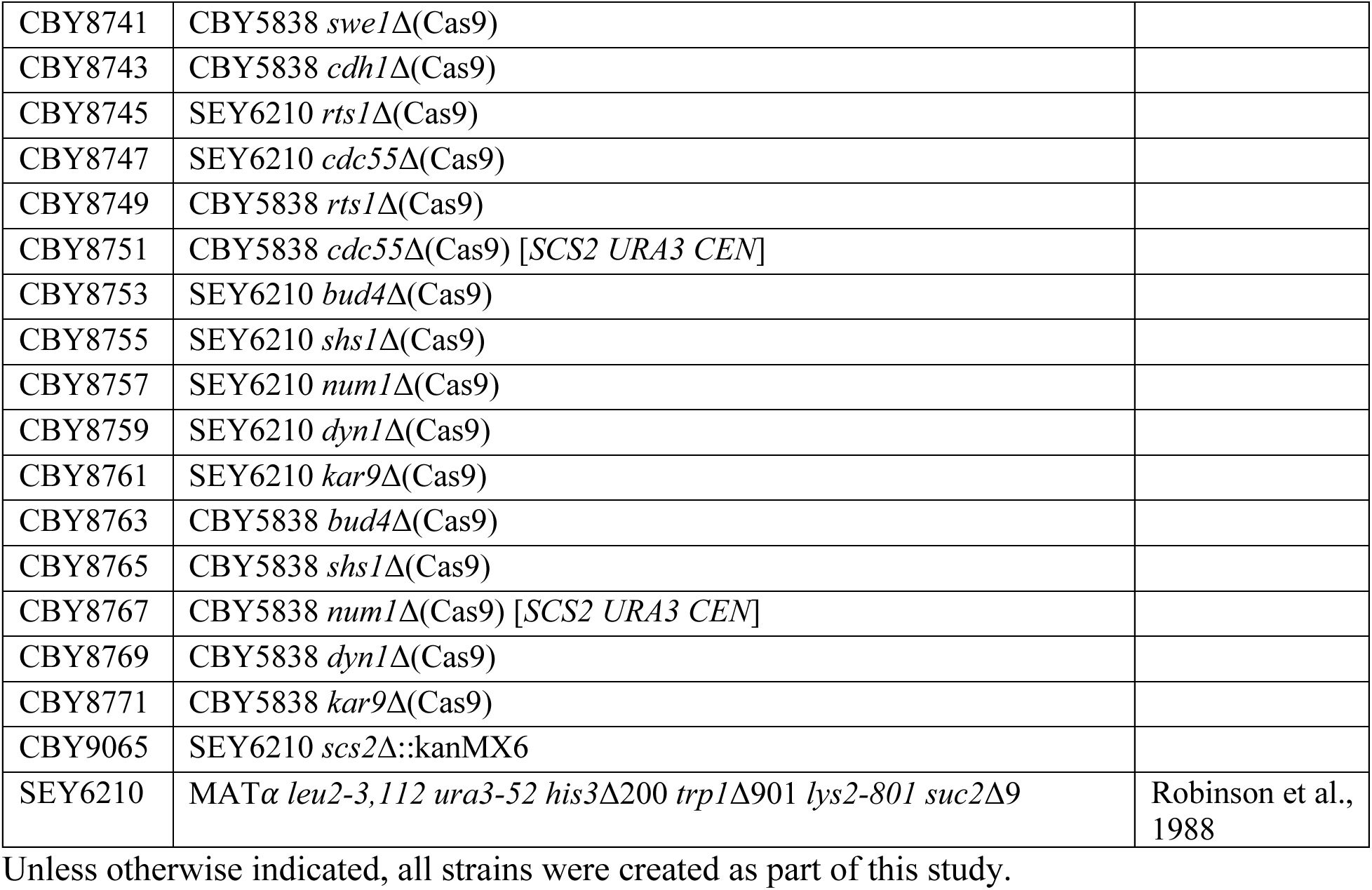
Yeast strains.

| **Strain** | **Genotype** | **Reference** |
|---|---|---|
| BY4741 | MAT**a** *leu2*Δ0 *ura3*Δ0 *his3*Δ0 *met15*Δ0 | Winzler et al., 1999 |
| BY4742 | MAT*α* *leu2*Δ0 *ura3*Δ0 *lys2*Δ0 *his3*Δ0 | Winzler et al., 1999 |
| CBY57 | MAT*α*/MAT**a** *leu2-3,112/leu2-3,112* *ura3-52/ura3-52* *his3Δ200/his3Δ200* *trp1*Δ901/ *trp1*Δ901 *suc2*Δ9/*suc2*Δ9 *lys2-801/LYS2* *ADE2*/*ade2*Δ |  |
| CBY5838 | SEY6210 *ist2*Δ::hisMX6 *scs2*Δ::TRP1 *scs22*Δ::hisMX6 *tcb1*Δ::kanMX6 *tcb2*Δ::kanMX6 *tcb3*Δ::hisMX6 *ice2*Δ::natMX4 | Quon et al., 2018 |
| CBY6095 | SEY6210 P<sup>*HIS3*</sup>-Venus-*TUB1*:hph |  |
| CBY6097 | SEY6210 P<sup>*HIS3*</sup>-mRuby2-*TUB1*:hph |  |
| CBY6107 | CBY5838 P<sup>*HIS3*</sup>-Venus-*TUB1*:hph |  |
| CBY6109 | CBY5838 P<sup>*HIS3*</sup>-mRuby2-*TUB1*:hph |  |
| CBY8089 | SEY6210 *CDC5*-GFP:*URA3* |  |
| CBY8091 | CBY5838 *CDC5*-GFP:*URA3* |  |
| CBY8597 | SEY6210 *rad52*Δ(Cas9) |  |
| CBY8695 | SEY6210 *snf7*Δ(Cas9) |  |
| CBY8697 | SEY6210 *chm7*Δ(Cas9) |  |
| CBY8699 | SEY6210 *heh1*Δ(Cas9) |  |
| CBY8701 | CBY5838 *heh1*Δ(Cas9) [*SCS2 URA3 CEN*] |  |
| CBY8703 | CBY5838 *chm7*Δ(Cas9) |  |
| CBY8705 | SEY6210 *nem1*Δ(Cas9) |  |
| CBY8707 | SEY6210 *pah1*Δ(Cas9) |  |
| CBY8709 | CBY5838 *nem1*Δ(Cas9) |  |
| CBY8711 | CBY5838 *pah1*Δ(Cas9) |  |
| CBY8713 | SEY6210 *chl4*Δ(Cas9) |  |
| CBY8715 | SEY6210 *bub3*Δ::hph |  |
| CBY8717 | CBY5838 *bub3*Δ::hph |  |
| CBY8719 | CBY5838 *chl4*Δ(Cas9) |  |
| CBY8721 | SEY6210 *bub2*Δ(Cas9) |  |
| CBY8723 | SEY6210 *kin4*Δ(Cas9) |  |
| CBY8725 | CBY5838 *bub2*Δ(Cas9) |  |
| CBY8727 | CBY5838 *kin4*Δ(Cas9) |  |
| CBY8729 | SEY6210 *elm1*Δ(Cas9) |  |
| CBY8731 | SEY6210 *hsl1*Δ(Cas9) |  |
| CBY8733 | SEY6210 *swe1*Δ(Cas9) |  |
| CBY8735 | SEY6210 *cdh1*Δ(Cas9) |  |
| CBY8737 | CBY5838 *elm1*Δ(Cas9) [*SCS2 URA3 CEN*] |  |
| CBY8739 | CBY5838 *hsl1*Δ(Cas9) |  |
| CBY8741 | CBY5838 *swe1*Δ(Cas9) |  |
| CBY8743 | CBY5838 *cdh1*Δ(Cas9) |  |
| CBY8745 | SEY6210 *rts1*Δ(Cas9) |  |
| CBY8747 | SEY6210 *cdc55*Δ(Cas9) |  |
| CBY8749 | CBY5838 *rts1*Δ(Cas9) |  |
| CBY8751 | CBY5838 *cdc55*Δ(Cas9) [*SCS2 URA3 CEN*] |  |
| CBY8753 | SEY6210 *bud4*Δ(Cas9) |  |
| CBY8755 | SEY6210 *shs1*Δ(Cas9) |  |
| CBY8757 | SEY6210 *num1*Δ(Cas9) |  |
| CBY8759 | SEY6210 *dyn1*Δ(Cas9) |  |
| CBY8761 | SEY6210 *kar9*Δ(Cas9) |  |
| CBY8763 | CBY5838 *bud4*Δ(Cas9) |  |
| CBY8765 | CBY5838 *shs1*Δ(Cas9) |  |
| CBY8767 | CBY5838 *num1*Δ(Cas9) [*SCS2 URA3 CEN*] |  |
| CBY8769 | CBY5838 *dyn1*Δ(Cas9) |  |
| CBY8771 | CBY5838 *kar9*Δ(Cas9) |  |
| CBY9065 | SEY6210 *scs2*Δ::kanMX6 |  |
| SEY6210 | MAT*α* *leu2-3,112* *ura3-52* *his3*Δ200 *trp1*Δ901 *lys2-801* *suc2*Δ9 | Robinson et al., 1988 |

Unless otherwise indicated, all strains were created as part of this study.

**S3 Table.**
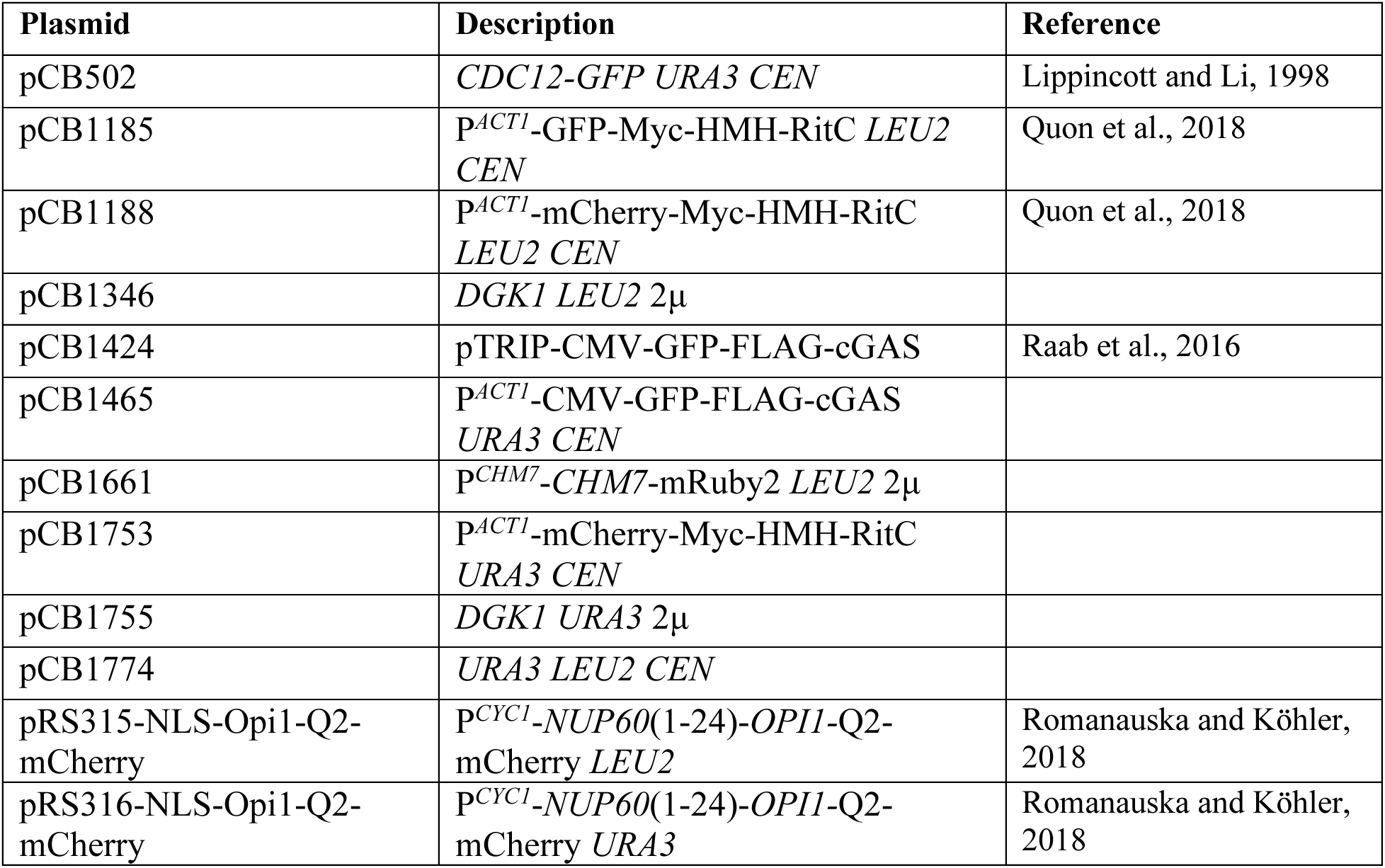
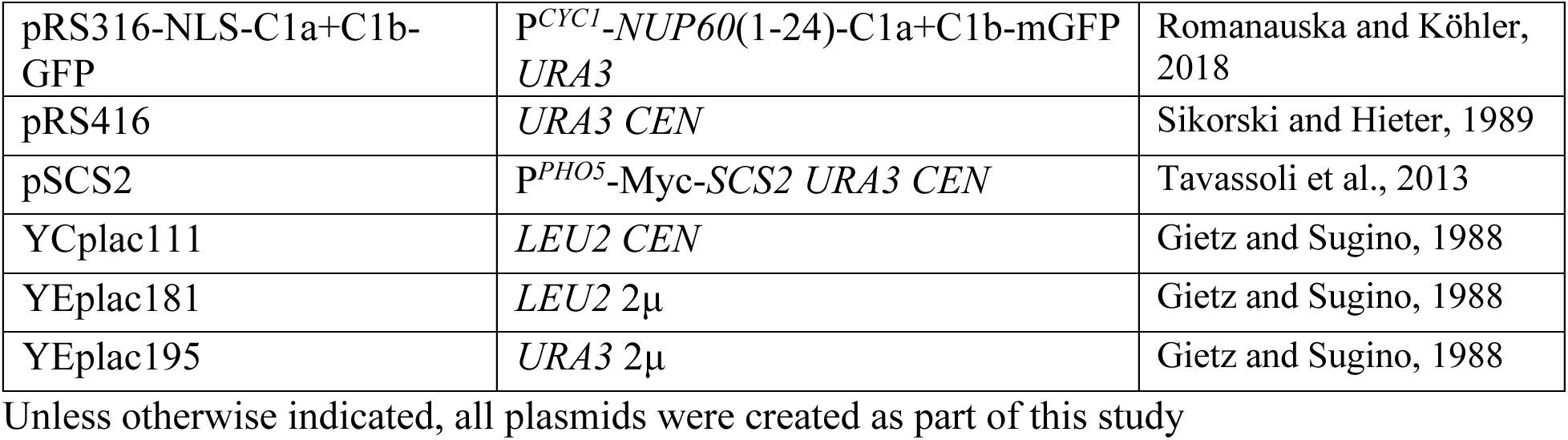
Plasmids.

**S1 Video. Time lapse of GFP-cGAS nuclear ruptures in Δ-s-tether cells during the cell cycle (corresponding to still frames shown in S1 Fig).** Example of unbudded Δ-s-tether cell (CBY5838) expressing GFP-cGAS (pCB1465) in which nuclear rupturing is first detected after 15 min; nuclear fluorescence persists as the daughter bud forms. The cell was imaged every 15 min for 1.5 hrs, and playback speed is 5 s = 90 min real time.

**S2 Video. Time lapse of highly motile intranuclear NLS-DAG-GFP aggregates.** Examples of WT (SEY6210) and Δ-s-tether (CBY5838) cells expressing the nuclear DAG probe (pRS316-NLS-C1a+C1b-GFP) cultured in liquid selective growth medium at 30°C. Highly motile intranuclear NLS-DAG-GFP particles vibrate within Δ-s-tether cells whereas comparable particles are not observed in WT. Nuclei (indicated by arrows) were identified by colocalization with the PA nuclear probe (not shown). WT cell image exposure was 200 ms for 10 s and Δ-s-tether cells were imaged every 100 ms for 10 s. The playback speed for all images corresponds to real time.

**S3 Video. Time lapse of microtubule-independent nucleopodial outgrowth in WT versus Δ-s-tether cells during preanaphase.** Examples of WT (CBY6095) and Δ-s-tether (CBY6107) cells expressing Venus-Tub1 and transformed with a plasmid expressing the nuclear NLS-PA-mCherry probe (pRS316-NLS-Opi1- Q2-mCherry). Microtubule-independent nucleopodia (red) are observed in most WT cells whereas only a minority of Δ-s-tether cells show nuclear membrane outgrowth that does not track with microtubules (yellow). The cell was imaged every 30 s for until the nucleus is midway through the bud neck, and playback speed is in real time.

**S1 Figure.**
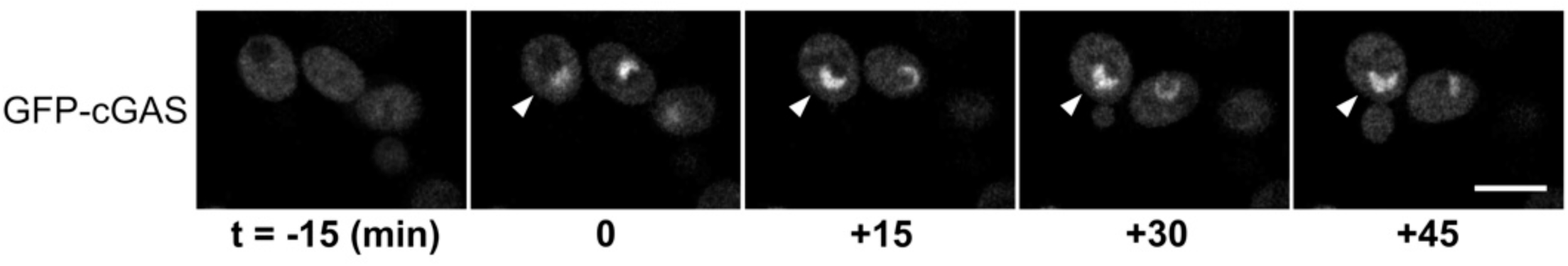
GFP-cGAS ruptures in Δ-s-tether cells during the cell cycle. Selected still frames corresponding to Video S1. The GFP-cGAS fluorescent nuclear rupture in the indicated unbudded Δ-s-tether (CBY5838) cell was apparent 15 min after the first image acquisition and fluorescence persisted during subsequent bud formation. GFP-cGAS nuclei are observed in budded and unbudded Δ-s-tether cells indicating that nuclear ruptures can occur throughout the cell cycle. Images were acquired at 15 min intervals.

**S2 Figure.**
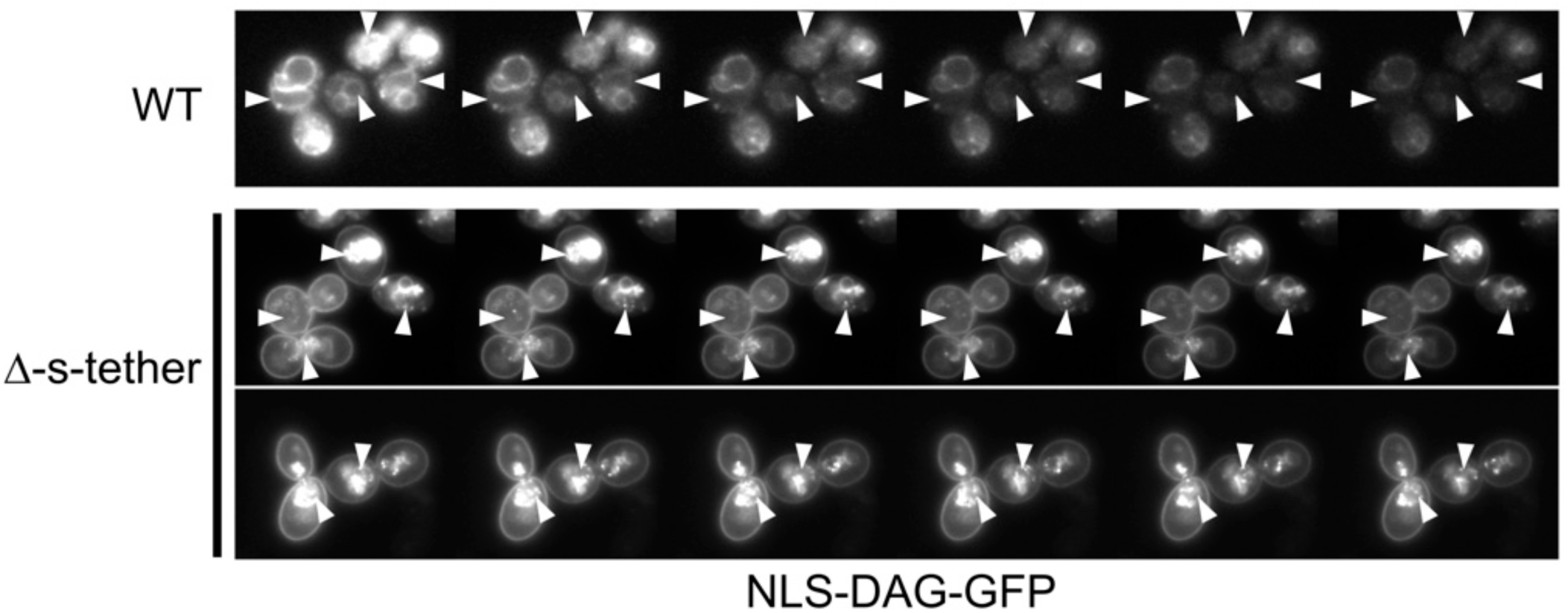
Still-frame image series of highly motile intranuclear NLS-DAG-GFP particles. Representative time-lapsed images of WT (SEY6210) and Δ-s-tether cells (CBY5838) expressing the nuclear DAG probe (pRS316-NLS-C1a+C1b-GFP). In contrast to Δ-s-tether cells, motile intranuclear NLS-DAG-GFP particles are not observed in WT. Nuclei were confirmed by colocalization with NLS-PA-mCherry. Images were acquired at 20 s intervals.

**S3 Figure.**
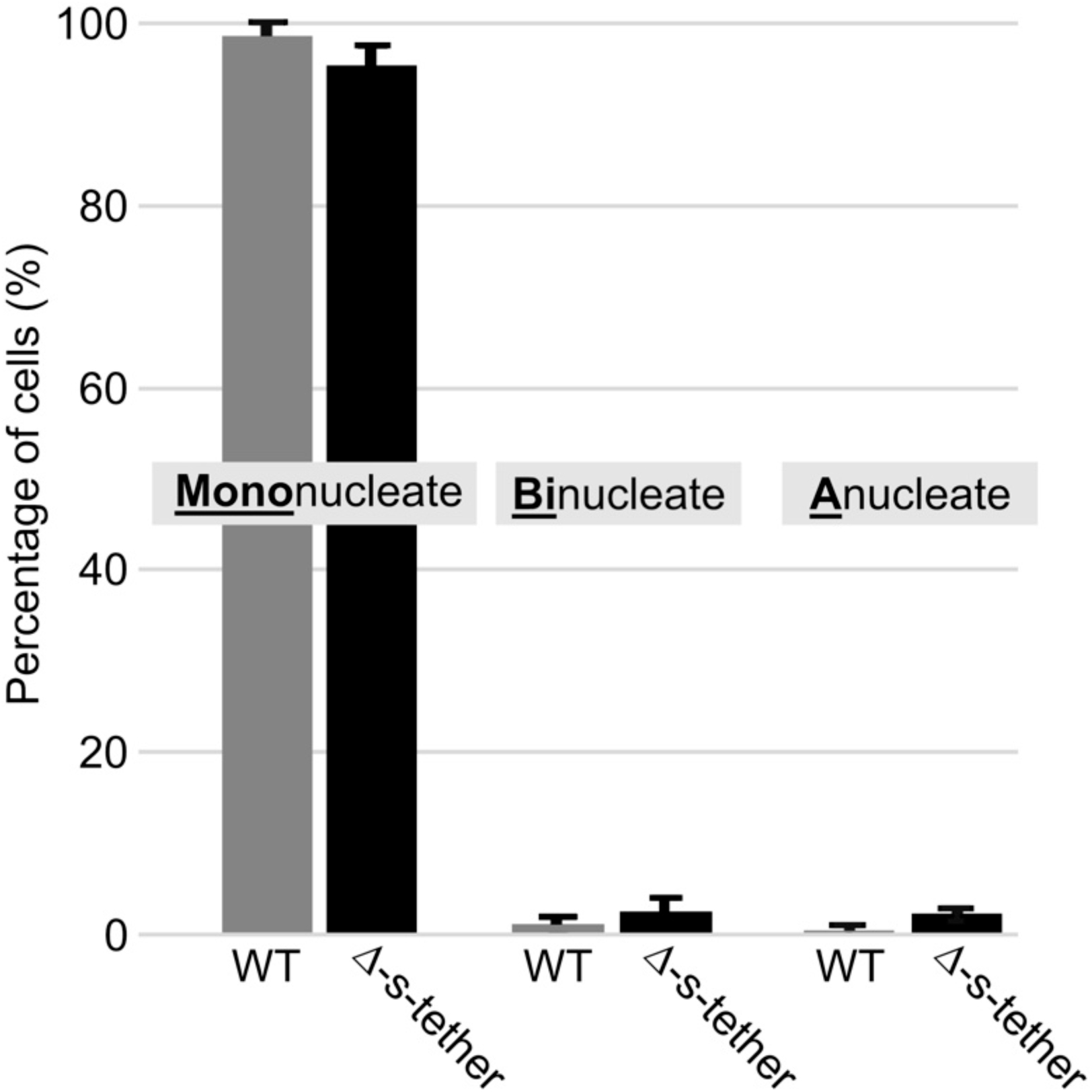
Quantification of binucleate Δ-s-tether cells versus WT. By DAPI nuclear fluorescence, the percentage of WT (SEY6210) and Δ-s-tether (CBY5838) cells that contain distinct nuclear DNA masses. Although Δ-s-tether cells exhibit a modestly higher percentage of binucleate and anucleate cells, the difference relative to WT is statistically insignificant (*p* > 0.1).

**S4 Figure.**
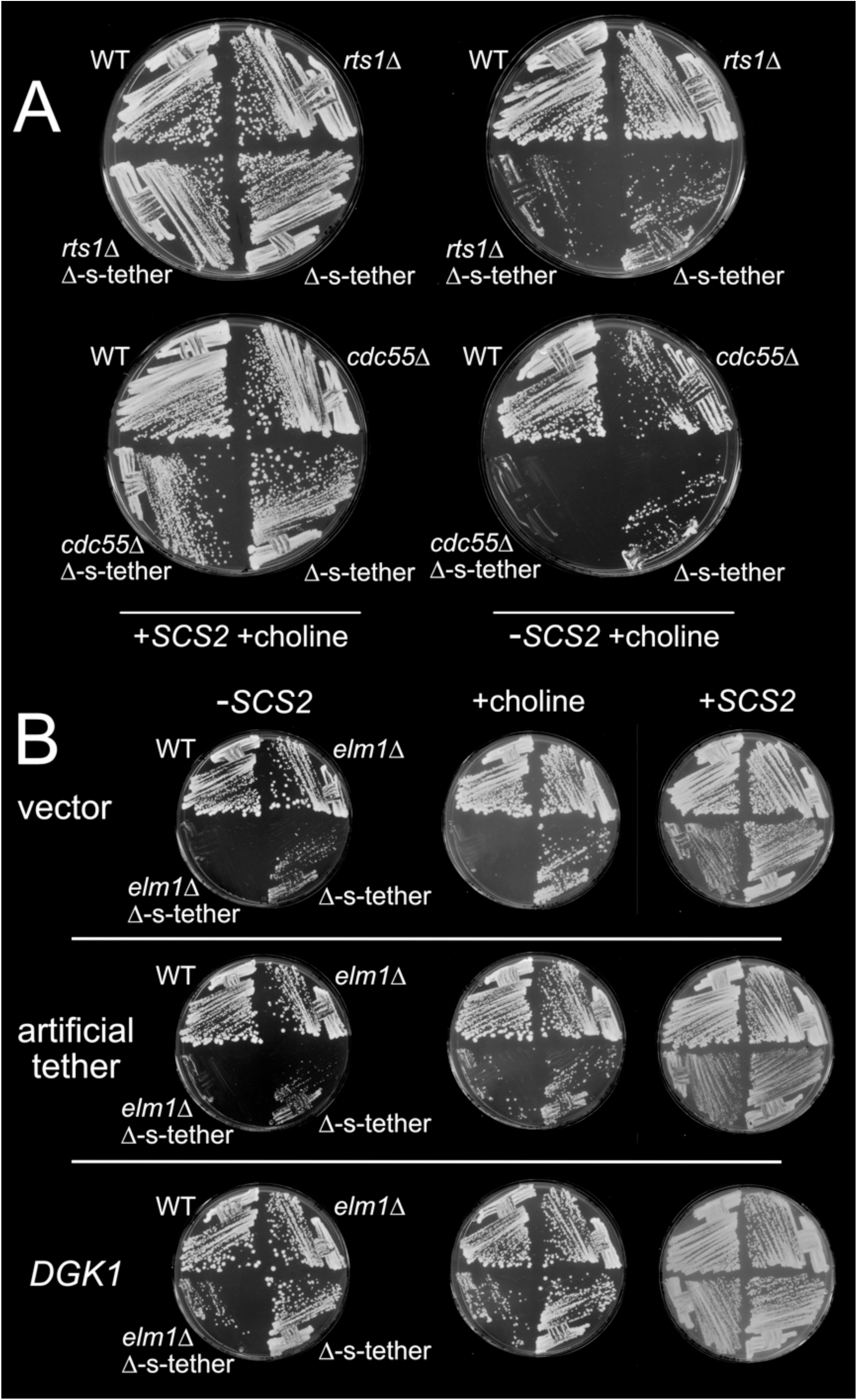
Lethality between Δ-s-tether and deletions of SPOC regulatory genes, and suppressor analysis. (A) WT (SEY6210), Δ-s-tether (CBY5838), *cdc55*Δ (CBY8747), *rts1*Δ (CBY8745), *cdc55*Δ Δ-s-tether (CBY8751), and *rts1*Δ Δ-s-tether (CBY8749) cells, with selection for (+*SCS2*) or against (-*SCS2*) the complementing *SCS2*-containing plasmid (pSCS2), streaked onto solid growth medium supplemented with 1 mM choline incubated for 4 days at 30°C. (B) WT, Δ-s-tether, *elm1*Δ (CBY8729), and *elm1*Δ Δ-s-tether (CBY8737) cells, selecting for (+*SCS2*) or against (-*SCS2*) the complementing *SCS2*-containing plasmid (pSCS2), streaked onto solid growth medium supplemented with 1 mM choline and incubated for 4 days at 30°C. As indicated, each strain was transformed with either a vector alone control (YEplac181), a plasmid expressing the artificial cER-PM staple (pCB1188), or a multicopy plasmid containing *DGK1* (pCB1346).

## Notes

### Competing Interest Statement

The authors have declared no competing interest.

### Funding

- Natural Sciences and Engineering Research Council of Canada: Discovery grant

## References

1. Rowat AC, Lammerding J, Herrmann H, Aebi U. Towards an integrated understanding of the structure and mechanics of the cell nucleus. Bioessays. 2008;30(3):226–236. doi:10.1002/bies.20720

2. Renvoisé B, Blackstone C. Emerging themes of ER organization in the development and maintenance of axons. Curr Opin Neurobiol. 2010;20(5):531–537. doi:10.1016/j.conb.2010.07.001

3. de Las Heras JI, Batrakou DG, Schirmer EC. Cancer biology and the nuclear envelope: a convoluted relationship. Semin Cancer Biol. 2013;23(2):125–137. doi:10.1016/j.semcancer.2012.01.008

4. Denais CM, Gilbert RM, Isermann P, McGregor AL, te Lindert M, Weiglin B, et al. Nuclear envelope rupture and repair during cancer cell migration. Science. 2016;352(6283):353–358. doi:10.1126/science.aad7297

5. Lim S, Quinton RJ, Ganem NJ. Nuclear envelope rupture drives genome instability in cancer. Mol Biol Cell. 2016;27(21):3210–3213. doi:10.1091/mbc.E16-02-0098

6. Arruda AP, Parlakgül G. Endoplasmic Reticulum Architecture and Inter-Organelle Communication in Metabolic Health and Disease. Cold Spring Harb Perspect Biol. 2023;15(2):a041261. doi:10.1101/cshperspect.a041261

7. Kuijpers M, Nguyen PT, Haucke V. The Endoplasmic Reticulum and Its Contacts: Emerging Roles in Axon Development, Neurotransmission, and Degeneration. Neuroscientist. 2024;30(5):545–559. doi:10.1177/10738584231162810

8. Du Y, Ferro-Novick S, Novick P. Dynamics and inheritance of the endoplasmic reticulum. J Cell Sci. 2004;117(Pt 14):2871–2878. doi:10.1242/jcs.01286

9. Manford AG, Stefan CJ, Yuan HL, Macgurn JA, Emr SD, et al. ER-to-plasma membrane tethering proteins regulate cell signaling and ER morphology. Dev Cell. 2012;23(6):1129–1140. doi:10.1016/j.devcel.2012.11.004

10. West M, Zurek N, Hoenger A, Voeltz GK. A 3D analysis of yeast ER structure reveals how ER domains are organized by membrane curvature. J Cell Biol. 2011;193(2):333–346. doi:10.1083/jcb.201011039

11. Henne WM, Liou J, Emr SD. Molecular mechanisms of inter-organelle ER-PM contact sites. Curr Opin Cell Biol. 2015;35:123–130. doi:10.1016/j.ceb.2015.05.001

12. Gallo A, Vannier C, Galli T. Endoplasmic Reticulum-Plasma Membrane associations: structures and functions. Annu Rev Cell Dev Biol. 2012;32:279–301. doi:10.1146/annurev-cellbio-111315-125024

13. Zaman MF, Nenadić A, Radojičić A, Rosado A, Beh CT. Sticking with it: ER-PM membrane contact sites as a coordinating nexus for regulating lipids and proteins at the cell cortex. Front Cell Dev Biol. 2020;8:675. doi:10.3389/fcell.2020.00675

14. Quon E, Sere YY, Chauhan N, Johansen J, Sullivan DP, Dittman JS, et al. Endoplasmic reticulum-plasma membrane contact sites integrate sterol and phospholipid regulation. PLoS Biol. 2018;16(5):e2003864. doi:10.1371/journal.pbio.2003864

15. Quon E, Nenadic A, Zaman MF, Johansen J, Beh CT. ER-PM membrane contact site regulation by yeast ORPs and membrane stress pathways. PLoS Genet. 2022;18(3):e1010106. doi:10.1371/journal.pgen.1010106

16. Nenadic A, Zaman MF, Johansen J, Volpiana MW, Beh CT. Increased Phospholipid Flux Bypasses Overlapping Essential Requirements for the Yeast Sac1p Phosphoinositide Phosphatase and ER-PM Membrane Contact Sites. J Biol Chem. 2023;299(9):105092. doi:10.1016/j.jbc.2023.105092

17. Winey M, Yarar D, Giddings TH Jr, Mastronarde DN. Nuclear pore complex number and distribution throughout the Saccharomyces cerevisiae cell cycle by three-dimensional reconstruction from electron micrographs of nuclear envelopes. Mol Biol Cell. 1997;8(11):2119–2132. doi:10.1091/mbc.8.11.2119

18. Santos-Rosa H, Leung J, Grimsey N, Peak-Chew S, Siniossoglou S. The yeast lipin Smp2 couples phospholipid biosynthesis to nuclear membrane growth. EMBO J. 2005;24(11):1931–1941. doi:10.1038/sj.emboj.7600672

19. Campbell JL, Lorenz A, Witkin KL, Hays T, Loidl J, Cohen-Fix O. Yeast nuclear envelope subdomains with distinct abilities to resist membrane expansion. Mol Biol Cell. 2006;17(4):1768–1778. doi:10.1091/mbc.e05-09-0839

20. Han GS, O’Hara L, Carman GM, Siniossoglou S. An unconventional diacylglycerol kinase that regulates phospholipid synthesis and nuclear membrane growth. J Biol Chem. 2008;283(29):20433–20442. doi:10.1074/jbc.M802903200

21. Siniossoglou S. Lipins, lipids and nuclear envelope structure. Traffic. 2009;10(9):1181–1187. doi:10.1111/j.1600-0854.2009.00923.x

22. Barbosa AD, Sembongi H, Su WM, Whittemore CM, Robenek H, Thiam AR, et al. Lipid partitioning at the nuclear envelope controls membrane biogenesis. Mol Biol Cell. 2015;26(20):3641–3657. doi:10.1091/mbc.E15-03-0173

23. Wang R, Kamgoue A, Normand C, Léger-Silvestre I, Mangeat T, Gadal O. High resolution microscopy reveals the nuclear shape of budding yeast during cell cycle and in various biological states. J Cell Sci. 2016;129(24):4480–4495. doi:10.1242/jcs.188250

24. Romanauska A, Köhler A. The inner nuclear membrane is a metabolically active territory that generates nuclear lipid droplets. Cell. 2018;174(3):700–715.e18. doi:10.1016/j.cell.2018.05.047

25. Zhang D, Oliferenko S. Remodeling the nuclear membrane during closed mitosis. Curr Opin Cell Biol. 2013;25(1):142–148. doi:10.1016/j.ceb.2012.09.001

26. Thaller DJ, Ader NR, Mannino M, Borah S, King MC, Ciani B, et al. Direct binding of ESCRT protein Chm7 to phosphatidic acid-rich membranes at nuclear envelope herniations. J Cell Biol. 2021;220(3):e202004222. doi:10.1083/jcb.202004222

27. Ptak C, Rehman S, Wozniak RW. Mechanisms of nuclear envelope expansion. Curr Opin Cell Biol. 2024;91:102425. doi:10.1016/j.ceb.2024.102425

28. Güttinger S, Laurell E, Kutay U. Orchestrating nuclear envelope disassembly and reassembly during mitosis. Nat Rev Mol Cell Biol. 2009;10(3):178–191. doi:10.1038/nrm2641

29. Boettcher B, Barral Y. The cell biology of open and closed mitosis. Nucleus. 2013;4(3):160–165. doi:10.4161/nucl.24676

30. Witkin WKL, Friederichs JM, Cohen-Fix O, Jaspersen SL. Changes in the nuclear envelope environment affect spindle pole body duplication in Saccharomyces cerevisiae. Genetics. 2010;186(3):867–883. doi:10.1534/genetics.110.119149

31. Meseroll RA, Cohen-Fix O. The malleable nature of the budding yeast nuclear envelope: Flares, fusion, and fenestrations. J Cell Physiol. 2016;231(11):2353–2360. doi:10.1002/jcp.25355

32. Bahmanyar S, Schlieker C. Lipid and protein dynamics that shape nuclear envelope identity. Mol Biol Cell. 2020;31(13):1315–1323. doi:10.1091/mbc.E18-10-0636

33. Foo S, Cazenave-Gassiot A, Wenk MR, Oliferenko S. Diacylglycerol at the inner nuclear membrane fuels nuclear envelope expansion in closed mitosis. J Cell Sci. 136(3):jcs260568. doi:10.1242/jcs.260568

34. Raab M, Gentili M, de Belly H, Thiam HR, Vargas P, Jimenez AJ, et al. ESCRT III repairs nuclear envelope ruptures during cell migration to limit DNA damage and cell death. Science. 2016;352(6283):359–362. doi:10.1126/science.aad7611

35. Warecki B, Ling X, Bast I, Sullivan W. ESCRT-III–mediated membrane fusion drives chromosome fragments through nuclear envelope channels. J Cell Biol. 2020;219(3):e201905091. doi:10.1083/jcb.201905091

36. Shah P, Wolf K, Lammerding J. Bursting the Bubble – Nuclear Envelope Rupture as a Path to Genomic Instability? Trends Cell Biol. 2017;27(8):546–555. doi:10.1016/j.tcb.2017.02.008

37. Kalukula Y, Stephens AD, Lammerding J, Gabriele S. Mechanics and functional consequences of nuclear deformations. Nat Rev Mol Cell Biol. 2022;23(9):583–602. doi:10.1038/s41580-022-00480-z

38. Olmos Y, Hodgson L, Mantell J, Verkade P, Carlton JG. ESCRT-III controls nuclear envelope reformation. Nature. 2015;522(7555):236–239. doi:10.1038/nature14503

39. Gu M, LaJoie D, Chen OS, von Appen A, Ladinsky MS, Redd MJ, et al. LEM2 recruits CHMP7 for ESCRT-mediated nuclear envelope closure in fission yeast and human cells. Proc Natl Acad Sci U S A. 2017;114(11):E2166–E2175. doi:10.1073/pnas.1613916114

40. Vietri M, Radulovic M, Stenmark H. The many functions of ESCRTs. Nat Rev Mol Cell Biol. 2020;21:25–42. doi:10.1038/s41580-019-0177-4

41. Kono Y, Shimi T. Crosstalk between mitotic reassembly and repair of the nuclear envelope. Nucleus. 2024;15(1):2352203. doi:10.1080/19491034.2024.2352203

42. Tang S, Henne WM, Borbat PP, et al. Structural basis for activation, assembly and membrane binding of ESCRT-III Snf7 filaments. Elife. 2015;4:e12548. doi:10.7554/eLife.12548

43. Gatta AT, Carlton JG. The ESCRT-machinery: Closing holes and expanding roles. Curr Opin Cell Biol. 2019;59:121–132. doi:10.1016/j.ceb.2019.04.005

44. Kirchenbauer D, Liakopoulos D. An auxiliary, membrane-based mechanism for nuclear migration in budding yeast. Mol Biol Cell. 2013;24(1):1–12. doi:10.1091/mbc.e12-08-0602

45. Xiang X. Nuclear movement in fungi. Semin Cell Dev Biol. 2018;82:3–16. doi:10.1016/j.semcdb.2017.10.024

46. Melloy PG, Rose MD. Influence of the bud neck on nuclear envelope fission in Saccharomyces cerevisiae. Exp Cell Res. 2017;358(2):390–396. doi:10.1016/j.yexcr.2017.07.013

47. Walters AD, May CK, Dauster ES, Cinquin BP, Smith EA, Robellet X, et al. The yeast polo kinase Cdc5 regulates the shape of the mitotic nucleus. Curr Biol. 2014;24(23):2861–2867. doi:10.1016/j.cub.2014.10.029

48. Botchkarev VV Jr, Haber JE. Functions and regulation of the Polo-like kinase Cdc5 in the absence and presence of DNA damage. Curr Genet. 2018;64(1):87–96. doi:10.1007/s00294-017-0727-2

49. Cid VCJ, Jiménez J, Molina MA, Sánchez M, Nombela C, Thorner JW. Orchestrating the cell cycle in yeast: sequential localization of key mitotic regulators at the spindle pole and the bud neck. Microbiology. 2002;148(Pt 9):2647–2659. doi:10.1099/00221287-148-9-2647

50. Park JE, Park CJ, Sakchaisri K, Karpova T, Asano S, McNally J, et al. Novel functional dissection of the localization-specific roles of budding yeast polo kinase Cdc5p. Mol Cell Biol. 2004;24(22):9873–9886. doi:10.1128/MCB.24.22.9873-9886.2004

51. Caydasi AK, Pereira G. SPOC alert--when chromosomes get the wrong direction. Exp Cell Res. 2012;318(12):1421–1427. doi:10.1016/j.yexcr.2012.03.031

52. Botchkarev VV Jr, Rossio V, Yoshida S. The budding yeast Polo-like kinase Cdc5 is released from the nucleus during anaphase for timely mitotic exit. Cell Cycle. 2014;13(20):3260–3270. doi:10.4161/15384101.2014.953882

53. Matellán L, Manzano-López J, Monje-Casas F. Polo-like kinase acts as a molecular timer that safeguards the asymmetric fate of spindle microtubule-organizing centers. eLife. 2020;9:e61488. doi:10.7554/eLife.61488

54. Estrada de Martin P, Du Y, Novick P, Ferro-Novick S. Ice2p is important for the distribution and structure of the cortical ER network in Saccharomyces cerevisiae. J Cell Sci. 2005;118(Pt 1):65–77. doi:10.1242/jcs.01583

55. Loewen CJ, Young BP, Tavassoli S, Levine TP. Inheritance of cortical ER in yeast is required for normal septin organization. J Cell Biol. 2007;179(3):467–483. doi:10.1083/jcb.200708205

56. Fehrenbacher J, Hyman AA, Minc N, Hagan I, Liakopoulos D. Nuclear positioning in yeast: the role of the cytoskeleton and the nuclear envelope. J Cell Biol. 2002;159(5):773–783. doi:10.1083/jcb.200207049

57. Bernales S, McDonald KL, Walter P. Autophagy counterbalances endoplasmic reticulum expansion during the unfolded protein response. PLoS Biol. 2006;4(12):e423. doi:10.1371/journal.pbio.0040423

58. Romanauska A, Köhler A. Lipid saturation controls nuclear envelope function. Nat Cell Biol. 2023;25:1290–1302. doi:10.1038/s41556-023-01207-8

59. Gordon LM, Sauerheber RD, Esgate JA, Dipple I, Marchmont RJ, Houslay MD. The increase in bilayer fluidity of rat liver plasma membranes achieved by the local anesthetic benzyl alcohol affects the activity of intrinsic membrane enzymes. J Biol Chem. 1980;255(10):4519–4527. doi:10.1016/0021-9258(80)80138-4

60. Schneiter R, Cole CN. Integrating complex functions: coordination of nuclear pore complex assembly and membrane expansion of the nuclear envelope requires a family of integral membrane proteins. Nucleus. 2010;1(5):387–392. doi:10.4161/nucl.1.5.12333

61. Fernandez-Martinez J, Phillips J, Sekedat MD, Diaz-Avalos R, Velazquez-Muriel J, et al. Structure-function mapping of a heptameric module in the nuclear pore complex. J Cell Biol. 2012;196(4):419–434. doi:10.1083/jcb.201109008

62. Vietri M, Schink KO, Campsteijn C, Wegner CS, Schultz SW, Christ L, et al. Spastin and ESCRT-III coordinate mitotic spindle disassembly and nuclear envelope sealing. Nature. 2015;522(7555):231–235. doi:10.1038/nature14408

63. Hurley JH. ESCRTs are everywhere. EMBO J. 2015;34(19):2398-2407. doi:10.15252/embj.201592484

64. Jorgensen JR, Tei R, Baskin JM, Michel AH, Kornmann B, Emr SD. ESCRT-III and ER-PM contacts maintain lipid homeostasis. Mol Biol Cell. 2020;31(12):1302–1313. doi:10.1091/mbc.E20-01-0061

65. Webster BM, Colombi P, Jäger J, Lusk CP. Surveillance of nuclear pore complex assembly by ESCRT- III/Vps4. Cell. 2014;159(2):388–401. doi:10.1016/j.cell.2014.09.012

66. Thaller DJ, Allegretti M, Borah S, Ronchi P, Beck M, et al. An ESCRT-LEM protein surveillance system is poised to directly monitor the nuclear envelope and nuclear transport system. eLife. 2019;8:e45284. doi:10.7554/eLife.45284

67. Sun L, Wu J, Du F, Chen X, Chen ZJ. Cyclic GMP-AMP synthase is a cytosolic DNA sensor that activates the type I interferon pathway. Science. 2013;339(6121):786–791. doi:10.1126/science.1232458

68. Decout A, Katz JD, Venkatraman S, et al. The cGAS–STING pathway as a therapeutic target in inflammatory diseases. Nat Rev Immunol. 2021;21:548–569. doi:10.1038/s41577-021-00524-z

69. Morita E, Colf LA, Karren MA, Sandrin V, Rodesch CK, et al. Human ESCRT-III and VPS4 proteins are required for centrosome and spindle maintenance. Proc Natl Acad Sci U S A. 2010;107(29):12889–12894. doi:10.1073/pnas.1005938107

70. Olmos Y, Perdrix-Rosell A, Carlton J. Membrane binding by CHMP7 coordinates ESCRT-III-dependent nuclear envelope reformation. Curr Biol. 2016;26(19):2635–2641. doi:10.1016/j.cub.2016.07.039

71. Gatta AT, Olmos Y, Stoten CL, Chen Q, Rosenthal PB, et al. CDK1 controls CHMP7-dependent nuclear envelope reformation. eLife. 2021;10:e59999. doi:10.7554/eLife.59999

72. Travers KJ, Patil CK, Wodicka L, Lockhart D, Weissman JS, Walter P. Functional and genomic analyses of response to unfolded protein in yeast protoplast. Cell. 2000;101(3):249–258. doi:10.1016/S0092-8674(00)80835-1

73. Gauthier BR, Comaills V. Nuclear envelope integrity in health and disease: Consequences on genome instability and inflammation. Int J Mol Sci. 22(14):7281. doi:10.3390/ijms22147281

74. Pascual F, Carman GM. Phosphatidate phosphatase, a key regulator of lipid homeostasis. Biochim Biophys Acta. 2013;1831(3):514–522. doi:10.1016/j.bbalip.2012.08.006

75. Karanasios E, Han GS, Xu Z, Carman GM, Siniossoglou S. A phosphorylation-regulated amphipathic helix controls the membrane translocation and function of the yeast phosphatidate phosphatase. Proc Natl Acad Sci U S A. 2010;107(41):17539–17544. doi:10.1073/pnas.1007974107

76. Maciejowski J, Hatch EM. Nuclear Membrane Rupture and Its Consequences. Annu Rev Cell Dev Biol. 2020;36:85–114. doi:10.1146/annurev-cellbio-020520-120627

77. Pfeifer CR, Xia Y, Zhu K, et al. Constricted migration increases DNA damage and independently represses cell cycle. Mol Biol Cell. 2018;29(16):1948–1962. doi:10.1091/mbc.E18-02-0079

78. Shah P, Hobson CM, Cheng S, et al. Nuclear deformation causes DNA damage by increasing replication stress. Curr Biol. 2021;31(4):753–765.e6. doi:10.1016/j.cub.2020.11.037

79. Xia Y, Ivanovska IL, Zhu K, et al. Nuclear rupture at sites of high curvature compromises retention of DNA repair factors. J Cell Biol. 2018;217(11):3796–3808. doi:10.1083/jcb.201711161

80. Xie G, Walker RR III, Irianto J. Nuclear mechanosensing: mechanism and consequences of a nuclear rupture. Mutat Res. 2020;821:111717. doi:10.1016/j.mrfmmm.2020.111717

81. Hervé S, Scelfo A, Bersano Marchisio G, et al. Chromosome mis-segregation triggers cell cycle arrest through a mechanosensitive nuclear envelope checkpoint. Nat Cell Biol. 2025;10.1038/s41556-024-01565-x. doi:10.1038/s41556-024-01565-x

82. Pot I, Measday V, Snydsman B, Cagney G, Fields S, Davis TN, et al. Chl4p and iml3p are two new members of the budding yeast outer kinetochore. Mol Biol Cell. 2003;14(2):460–476. doi:10.1091/mbc.e02-08-0517

83. Medina-Suárez S, Machín F. Continuous nuclear envelope surveillance is required for DNA double strand break repair. Commun Biol. 2025;8(1):984. doi:10.1038/s42003-025-08416-x

84. Lang GI, Murray AW. Estimating the per-base-pair mutation rate in the yeast Saccharomyces cerevisiae. Genetics. 2008;178(1):67–82. doi:10.1534/genetics.107.071506

85. Simon DN, Wilson KL. The nucleoskeleton as a genome-associated dynamic ’network of networks’. Nat Rev Mol Cell Biol. 2011;12(11):695–708. doi:10.1038/nrm3207

86. Liakopoulos D, Kusch J, Grava S, Vogel J, Barral Y. Asymmetric loading of Kar9 onto spindle poles and microtubules ensures proper spindle alignment. Cell. 2003;112(4):561–574. doi:10.1016/S0092-8674(03)00119-3

87. Orellana-Muñoz S, Dueñas-Santero E, Arnaíz-Pita Y, Del Rey F, Correa-Bordes J, Vazquez de Aldana CR. The anillin-related Int1 protein and the Sep7 septin collaborate to maintain cellular ploidy in Candida albicans. Sci Rep. 2018;8(1):2257. doi:10.1038/s41598-018-20249-9

88. Lew DJ, Burke DL. The spindle assembly and spindle position checkpoints. Annu Rev Genet. 2003;37:251–282. doi:10.1146/annurev.genet.37.042203.120656

89. Li G, Moore JK. Microtubule dynamics at low temperature: evidence that tubulin recycling limits assembly. Mol Biol Cell. 2020;31(11):1154–1166. doi:10.1091/mbc.E19-11-0634

90. Bouquin N, Barral Y, Courbeyrette R, Blondel M, Snyder M, Mann C. Regulation of cytokinesis by the Elm1 protein kinase in Saccharomyces cerevisiae. J Cell Sci. 2000;113(Pt 8):1435–1445. doi:10.1242/jcs.113.8.1435

91. Thomas CL, Blacketer MJ, Edgington NP, Myers AM. Assembly interdependence among the S. cerevisiae bud neck ring proteins Elm1p, Hsl1p and Cdc12p. Yeast. 2003;20(9):813-826. doi:10.1002/yea.1003

92. Caydasi AK, Kurtulmus B, Orrico MI, Hofmann A, Ibrahim B, Pereira G. Elm1 kinase activates the spindle position checkpoint kinase Kin4. J Cell Biol. 2010;190(6):975–989. doi:10.1083/jcb.201006151

93. Perez AM, Finnigan GC, Roelants FM, Thorner J. Septin-Associated Protein Kinases in the Yeast Saccharomyces cerevisiae. Front Cell Dev Biol. 2016;4:119. doi:10.3389/fcell.2016.00119

94. Marquardt J, Chen X, Bi E. Reciprocal regulation by Elm1 and Gin4 controls septin hourglass assembly and remodeling. J Cell Biol. 2024;223(5):e202308143. doi:10.1083/jcb.202308143

95. Moyano-Rodríguez Y, Queralt E. PP2A Functions during Mitosis and Cytokinesis in Yeasts. Int J Mol Sci. 2020;21:264. doi:10.3390/ijms21010264

96. Chan LY, Amon A. The protein phosphatase 2A functions in the spindle position checkpoint by regulating the checkpoint kinase Kin4. Genes Dev. 2009;23(14):1639–1649. doi:10.1101/gad.1804609

97. Harvey SL, Enciso G, Dephoure N, Liakopoulos D, Kellogg DR. A phosphatase threshold sets the level of Cdk1 activity in early mitosis in budding yeast. Mol Biol Cell. 2011;22(19):3595–3608. doi:10.1091/mbc.E11-04-0340

98. Baró B, Játiva S, Calabria I, Vinaixa J, Bech-Serra JJ, de La Torre C, et al. SILAC-based phosphoproteomics reveals new PP2A-Cdc55-regulated processes in budding yeast. GigaScience. 2018;7(5):giy047. doi:10.1093/gigascience/giy047

99. Hu F, Wang Y, Liu D, Li Y, Qin J, Elledge SJ. Regulation of the Bub2/Bfa1 GAP complex by Cdc5 and cell cycle checkpoints. Cell. 2001;107(5):655–665. doi:10.1016/S0092-8674(01)00580-3

100. Geymonat M, Glasspar ML, Jesch J, et al. In vitro regulation of budding yeast Bfa1/Bub2 GAP activity by Cdc5. J Biol Chem. 2003;278(17):14591–14594. doi:10.1074/jbc.C300059200

101. Moore JK, Chudalayandi P, Heil-Chapdelaine RA, Cooper JA. The spindle position checkpoint is coordinated by the Elm1 kinase. J Cell Biol. 2010;191(3):493–503. doi:10.1083/jcb.201006092

102. Visintin R, Hwang E, Amon A. Regulation of the mitotic exit in Saccharomyces cerevisiae. Science. 1898;282(5399):1034–1037. doi:10.1126/science.282.5399.1034

103. Jaspersen SL, Goh PY, Winey M, Emr SD. Molecular analysis of the role of Cdc14p in the regulation of the anaphase-promoting complex in Saccharomyces cerevisiae. J Cell Biol. 1999;144(6):1241–1250. doi:10.1083/jcb.144.6.1241

104. Cohen-Fix O, Peters JM, Kirschner MW. The role of Cdc14p in the regulation of the cell cycle in Saccharomyces cerevisiae. J Cell Biol. 2000;149(3):679–688. doi:10.1083/jcb.149.3.679

105. Shirayama M, Zachariae W, Ciosk R, Nasmyth K. The Polo-like kinase Cdc5p and the WD-repeat protein Cdc20p/fizzy are regulators and substrates of the anaphase promoting complex in Saccharomyces cerevisiae. EMBO J. 1998;17(5):1336–1349. doi:10.1093/emboj/17.5.1336

106. Rahal R, Amon A. The Polo-like kinase Cdc5 interacts with FEAR network components and Cdc14. Cell Cycle. 2008;7(20):6852–6860. doi:10.4161/cc.7.20.6852

107. Visintin C, Tomson BN, Rahal R, Paulson J, Cohen M, et al. APC/C-Cdh1-mediated degradation of the Polo kinase Cdc5 promotes the return of Cdc14 into the nucleolus. Genes Dev. 2008;22(1):79–90. doi:10.1101/gad.1601308

108. Lahiri S, Toulmay A, Prinz WA. Membrane contact sites, gateways for lipid homeostasis. Curr Opin Cell Biol. 2015;33:82–87. doi:10.1016/j.ceb.2014.12.004

109. Hatch EM. Nuclear envelope rupture: little holes, big openings. Curr Biol. 2018;52:66–72. doi:10.1016/j.cub.2018.02.001

110. Kucińska MK, Molinari M. Control of nuclear envelope dynamics during acute ER stress by LINC complexes disassembly and selective, asymmetric autophagy of the outer nuclear membrane. Autophagy. 2024;20(5):1194–1196. doi:10.1080/15548627.2023.2299123

111. Chen Z, Chen P, Li J, Landao-Bassonga E, Papadimitriou J, et al. External strain on the plasma membrane is relayed to the endoplasmic reticulum by membrane contact sites and alters cellular energetics. Sci Adv. 2025;11(26):eads6132. doi:10.1126/sciadv.ads6132

112. Lin F, Blake DL, Callebaut I, Skerjanc IS, Holmer L, et al. MAN1, an inner nuclear membrane protein that shares the LEM domain with lamina-associated polypeptide 2 and emerin. J Biol Chem. 2000;275(7):4840–4847. doi:10.1074/jbc.275.7.4840

113. Gonzalez Y, Saito A, Sazer S. Fission yeast Lem2 and Man1 perform fundamental functions of the animal cell nuclear lamina. Nucleus. 2012;3(1):60–76. doi:10.4161/nucl.18824

114. Taddei A, Gasser SM. Structure and function in the budding yeast nucleus. Genetics. 2012;192(1):107–129. doi:10.1534/genetics.112.140608

115. Odell J, Lammerding J. Lamins as structural nuclear elements through evolution. Curr Opin Cell Biol. 2023;85:102267. doi:10.1016/j.ceb.2023.102267

116. Houben F, Ramaekers FC, Snoeckx LH, Broers JL. Role of nuclear lamina-cytoskeleton interactions in the maintenance of cellular strength. Biochim Biophys Acta. 2007;1773(5):675–686. doi:10.1016/j.bbamcr.2006.09.018

117. Turgay Y, Eibauer M, Goldman AE, Shimi T, Khayat M, et al. The molecular architecture of lamins in somatic cells. Nature. 2017;543(7644):261–264. doi:10.1038/nature21382

118. McGillivary RM, Starr DA, Luxton GWG. Building and breaking mechanical bridges between the nucleus and cytoskeleton: Regulation of LINC complex assembly and disassembly. Curr Opin Cell Biol. 2023;85:102260. doi:10.1016/j.ceb.2023.102260

119. Hattier T, Andrulis ED, Tartakoff AM. Immobility, inheritance and plasticity of shape of the yeast nucleus. BMC Cell Biol. 2007;8:47. doi:10.1186/1471-2121-8-47

120. Fan J, Sun Z, Wang Y. The assembly of a noncanonical LINC complex in Saccharomyces cerevisiae. Curr Genet. 2022;68(1):91–96. doi:10.1007/s00294-021-01220-0

121. Wu Y, Whiteus C, Xu CS, Hayworth KJ, Weinberg RJ, Hess HF, et al. Contacts between the endoplasmic reticulum and other membranes in neurons. Proc Natl Acad Sci U S A. 2017;114(24):E4859–E4867. doi:10.1073/pnas.1701078114

122. Chen YJ, Quintanilla CG, Liou J. Recent insights into mammalian ER-PM junctions. Curr Opin Cell Biol. 2019;57:99–105. doi:10.1016/j.ceb.2018.12.011

123. Hwang S, Russo W, Cormier J, Johnson J, Martin S, Ippolito MR, et al. Sphingolipid synthesis maintains nuclear membrane integrity and genome stability during cell division. J Cell Biol. 2025;224(8):e202407209. doi:10.1083/jcb.202407209

124. Lindahl T, Barnes DE, Yang YG, Robins P. Biochemical properties of mammalian TREX1 and its association with DNA replication and inherited inflammatory disease. Biochem Soc Trans. 2009;37(Pt 3):535–538. doi:10.1042/BST0370535

125. Maciejowski J, Li Y, Bosco N, Campbell PJ, de Lange T. Chromothripsis and kataegis induced by telomere crisis. Cell. 2015;163(7):1641–1654. doi:10.1016/j.cell.2015.11.054

126. Krupina K, Goginashvili A, Baughn MW, et al. Chromothripsis and ecDNA initiated by N4BP2 nuclease fragmentation of cytoplasm-exposed chromosomes. Science. 2025;390(6778):1156–1163. doi:10.1126/science.ado0977

127. Nishikawa S, Terazawa Y, Nakayama T, Hirata A, Makio T, Endo T. Nep98p is a component of the yeast spindle pole body and essential for nuclear division and fusion. J Biol Chem. 2003;278(11):9938–9943. doi:10.1074/jbc.M210934200

128. Juanes MA, Piatti S. The final cut: cell polarity meets cytokinesis at the bud neck in S. cerevisiae. Cell Mol Life Sci. 2016;73(16):3115–3136. doi:10.1007/s00018-016-2220-3

129. Akhmetov A, Laurent JM, Gollihar J, Gardner EC, Garge RK, Ellington AD, et al. Single-step precision genome editing in yeast using CRISPR-Cas9. Bio-protocol. 2018;8(6):e2765. doi:10.21769/BioProtoc.2765

130. Lee ME, DeLoache WC, Cervantes B, Dueber JE. A highly characterized yeast toolkit for modular, multipart assembly. ACS Synth Biol. 2015;4(9):975–986. doi:10.1021/sb500366v

131. Todd RT, Braverman A, Selmecki A. Flow cytometry analysis of fungal ploidy. Curr Protoc Microbiol. 2018;50(1):e58. doi:10.1002/cpmc.58

132. Alfaro G, Johansen J, Dighe SA, Duamel G, Kozminski KG, Beh CT. The sterol-binding protein Kes1/Osh4p is a regulator of polarized exocytosis. Traffic. 2011;12(11):1521–1536. doi:10.1111/j.1600-0854.2011.01265.x

133. Kozminski KG, Alfaro G, Dighe S, Beh CT. Homologues of oxysterol-binding proteins affect Cdc42p- and Rho1p-mediated cell polarization in Saccharomyces cerevisiae. Traffic. 2006;7(9):1224–1242. doi:10.1111/j.1600-0854.2006.00467.x

134. Liachko I, Dunham MJ. An autonomously replicating sequence for use in a wide range of budding yeasts. FEMS Yeast Res. 2014;14(2):364–367. doi:10.1111/1567-1364.12123

135. Metzger MB, Scales JL, Dunklebarger MF, Weissman AM. The Ubiquitin Ligase (E3) Psh1p Is Required for Proper Segregation of both Centromeric and Two-Micron Plasmids in Saccharomyces cerevisiae. G3 (Bethesda). 2017;7(11):3731–3743. doi:10.1534/g3.117.300227

136. Zheng Q. Methods for comparing mutation rates using fluctuation assay data. Mutat Res. 2015;777:20–22. doi:10.1016/j.mrfmmm.2015.04.002

137. Zheng Q. rSalvador: An R package for the fluctuation experiment. G3 (Bethesda). 2017;7(12):3849–3856. doi:10.1534/g3.117.300120

